# Inter-organ Wingless/Ror/Akt signaling regulates nutrient-dependent hyperarborization of somatosensory neurons

**DOI:** 10.1101/2022.04.29.489983

**Authors:** Yasutetsu Kanaoka, Koun Onodera, Kaori Watanabe, Tadao Usui, Tadashi Uemura, Yukako Hattori

## Abstract

Nutrition in early life has profound effects on an organism, altering processes such as organogenesis. However, little is known about how specific nutrients affect neuronal development. Dendrites of class IV neurons in *Drosophila* larvae become more complex when the larvae are reared on a low-yeast diet compared to a high-yeast diet. Our systematic search for key nutrients revealed that the neurons increase their dendrite densities in response to a combined deficiency in vitamins, metal ions, and cholesterol. These nutrients affect dendrite branching through the Wingless/Ror/Akt pathway between the neuron and closely located muscles, and this short range-pathway is regulated by a systemic Upd2-Stat92E pathway between the fat body and muscles. Additionally, the low-yeast diet blunts neuronal light responsiveness and light avoidance behavior, which may help larvae optimize their survival strategies under low-nutritional conditions. Together, our studies illustrate how the availability of specific nutrients affects neuronal development through inter-organ signaling.

## INTRODUCTION

The physiological state of an organism influences organogenesis throughout the body. Among many external factors affecting the physiological state, nutrition in early life has a profound impact (Bhutta et al., 2017). This is particularly the case with neural development, which is highly metabolically demanding. A large amount of energy is consumed to control neural stem cell division, form complex dendrites and long axons in myriad neuronal cell types, and ultimately construct functional neural circuits (Prado and Dewey, 2014; Georgieff et al., 2015). Compared to metabolic regulation of neural stem cell proliferation (Homem et al., 2015), little is known about how nutritional status is conveyed to developing neurons and how those neurons regulate growth in response to such a signal (Shimada-Niwa and Niwa 2014; Shimono et al., 2014; Liu et al., 2017).

Dietary nutrients are absorbed by the digestive tract and circulated throughout the body, and they are sensed by organs including the nervous system (Chantranupong et al., 2015). Those organs communicate the nutritional status to each other by secreting signaling molecules, either low-molecular weight metabolites or macromolecules such as soluble proteins and lipoprotein particles, to elicit tissue-specific responses; and it is this inter-organ communication network that coordinates organogenesis with body growth (Droujinine and Perrimon 2016; Texada et al., 2020). In the nervous system, neurons sense circulating nutrients directly or by way of signaling molecules derived from other tissues, so there exist diverse modes of nutrient sensing (Morton et al., 2014; Jayakumar and Hasan, 2018).

The above-mentioned regulatory mechanisms of nutrient-dependent neuronal development can be explored at the molecular level with appropriate model neurons; and one particularly amenable model is the *Drosophila* class IV dendritic arborization neuron located in the larval periphery (class IV neurons hereafter; Grueber et al., 2002). Dendritic arbors of class IV neurons extensively cover the body wall, and they are elaborated two-dimensionally between the epidermis and the body wall muscles. Class IV neurons in larvae respond to noxious thermal, mechanical, and light stimuli and provoke robust avoidance behaviors (Tracey et al., 2003; Hwang et al., 2007; Xiang et al., 2010; Zhong et al., 2010; Guntur et al., 2015; Chin and Tracey, 2017). In the context of adaptation of growing animals to nutritional environments, it has been shown more recently that the class IV neurons sense amino acid deprivation by an amino acid transporter, Slimfast, at a late larval stage, which contributes to overcoming the nutritional stress, thereby allowing pupariation (Jayakumar et al., 2016, 2018). In addition, we and another group have shown that dendrites of class IV neurons become more complex when larvae are reared on low-yeast diets compared to high-yeast diets (Figures 1A, 1B, and Figure 1—figure supplement 1A; Watanabe et al., 2017; Poe et al., 2020). We designate this counterintuitive phenotype as hyperarborization. Although the entire larval development takes longer on the low-yeast diet compared to the high-yeast diet (Figure 1—figure supplement 1A), it is unlikely that the hyperarborization is a simple consequence of the longer larval stage. This is because a high-sugar diet also extends larval development, but the hyperarborization is not observed (Watanabe et al., 2017). Therefore, it has been assumed that the low-yeast diet is deficient in select nutrients, which causes the phenotype. However, the identities of such key nutrients responsible for the hyperarborization phenotype have heretofore not been determined.

**Figure 1.**
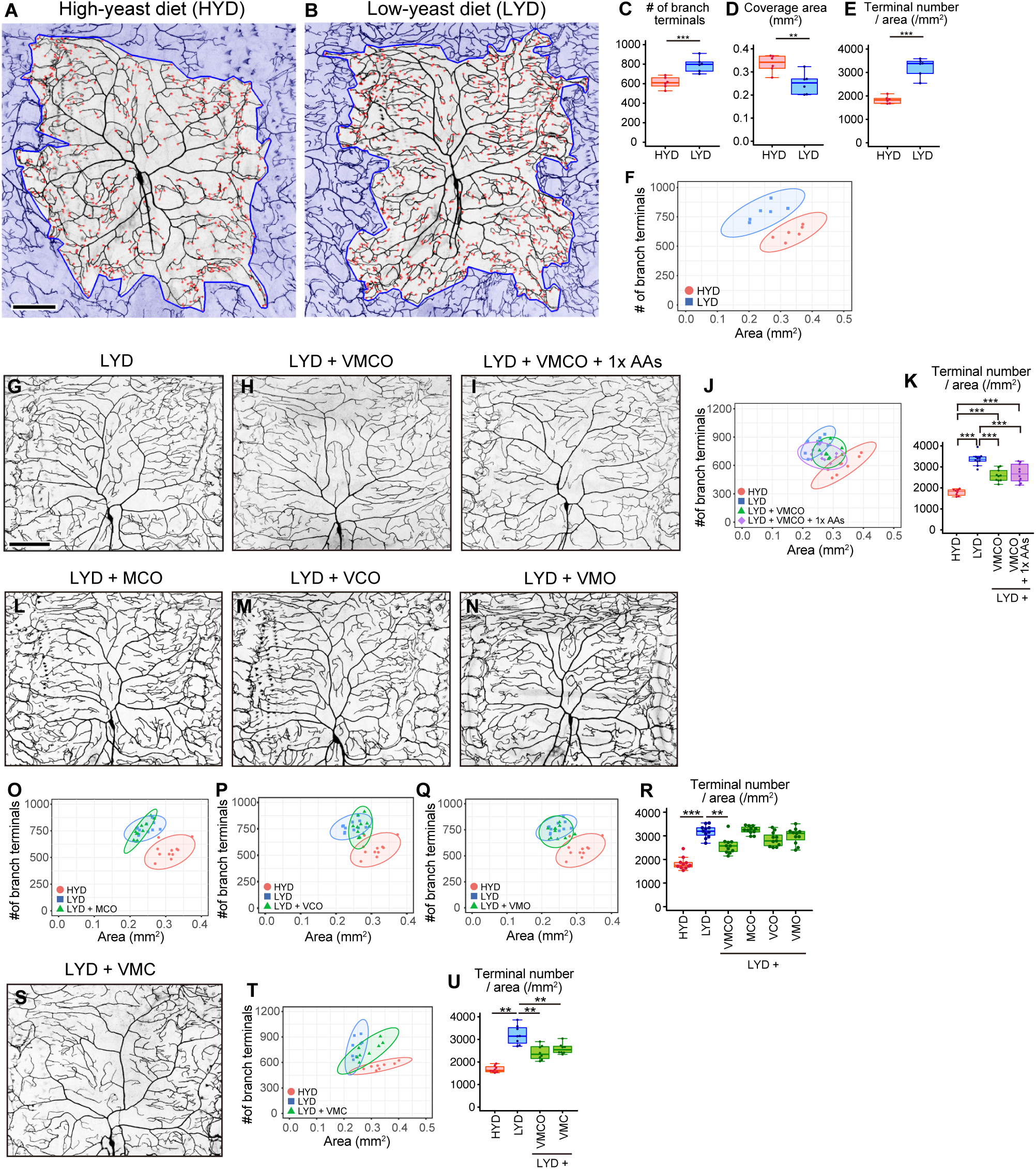
A mixture of vitamins, metal ions, and cholesterol ameliorates class IV neuron hyperarborization. (A and B) Representative output images of DeTerm. DeTerm automatically detects dendritic terminals of class IV neuron ddaC in larvae reared on a high-yeast diet (HYD; A) or a low-yeast diet (LYD; B). Red points indicate detected branch terminals. (C-E) The numbers of branch terminals detected by DeTerm (C), coverage areas of dendrites (D), and densities of branch terminals (the number of branch terminals/coverage area; E) of individual neurons, on HYD or LYD (Student’s t-test, n=6). Boxes show the 25th–75th percentiles. The central lines indicate the medians. Whiskers extend to the most extreme data points, which are no more than 1.5 times the interquartile range. Boxes and points for HYD data and those for LYD data are colored red and blue, respectively, in this and subsequent figures. (F) Two-dimensional plot of the dendritic area and the number of branch terminals of each neuron. The ellipses represent the 95% confidence intervals, which are clearly separated for HYD and LYD. (G-I) Images of ddaC neurons in larvae reared on LYD (G), LYD + vitamin + metal ion + cholesterol + other ingredients (LYD + VMCO; H), or LYD + VMCO + 1x amino acids (LYD + VMCO + 1x AAs; I). (J and K) Quantitative analysis of ddaC on LYD + VMCO or LYD + VMCO + 1x AAs. (J) Two-dimensional plot. Note that the ellipse of LYD +VMCO and that of LYD + VMCO + 1x AAs are located between those of HYD and LYD. (K) Densities of branch terminals (One-way ANOVA and Tukey’s HSD test, n = 8-10). (L-N) Images of ddaC neurons in larvae reared on LYD + metal ion + cholesterol + other ingredients (LYD + MCO; L), LYD + vitamin + cholesterol + other ingredients (LYD + VCO; M), or LYD + vitamin + metal ion + other ingredients (LYD + VMO; N). (O-R) Two-dimensional plots of ddaC on LYD + MCO (O), LYD + VCO (P), or LYD + VMO (Q), and densities of branch terminals (Steel test, n = 10; R). The elipses of these diets largely overlap with that of LYD and clearly or almost separare from that of HYD (O-Q). (S) Images of ddaC neurons in larvae reared on LYD + vitamin + metal ion + cholesterol (LYD + VMC) (T and U) Two-dimensional plots of ddaC neurons in larvae reared on LYD + VMC (T) and densities of branch terminals (Steel test, n = 8; U). The ellipse of LYD +VMC is located between those of HYD and LYD (T). Boxplots in (K, R and U) are depicted as in Figure 1C. *P < 0.05, **P < 0.01, and ***P < 0.001. Scale bars, 100μm. Figure supplement for figure 1: Figure supplement 1. Addition of amino acids does not rescue the hyperarborization

A wealth of genetic analyses on standard foods has revealed numerous regulators of dendrite morphogenesis working either in cell-autonomous or non-cell autonomous manners (Jan and Jan, 2010; Dong et al., 2015; Valnegri et al., 2015). Some of the cell-autonomous mechanisms include those related to intake and synthesis of metabolites: amino acid transporter SLC36/Pathetic (Path) (Lin et al., 2015) and a critical regulator of fatty acid synthesis, sterol regulatory element binding protein (SREBP) (Meltzer et al., 2017; Ziegler et al., 2017). Concerning the non-cell autonomous mechanisms, direct interactions between class IV neurons and one of the adjacent tissues, the epidermis, have been well characterized with the help of anatomical approaches under both light and electron microscopes (Yang and Chien, 2019). Some portions of dendritic branches are attached to the extracellular matrix, and the attachment is mediated by signaling between an epidermally-derived semaphoring ligand Sema-2b and its receptor Plexin B (PlexB) on the dendrite (Meltzer et al., 2016), as well as between a TGF-β ligand Maverick (Mav)-Ret receptor combination (Hoyer et al., 2018). Other portions of dendritic arbors are wrapped by epidermal cells, so overall the dendrite arbor is embedded in the epidermis locally (Han et al., 2012; Kim et al., 2012; Tenenbaum et al., 2017; Jiang et al., 2019). In contrast to the above dendrite-epidermis interaction, there is much less evidence for signaling between muscles and dendrites, despite their proximity to dendrites and their large volume in the body (Yasunaga et al., 2010). Furthermore, when considering the relationship between nutritional status and class IV neurons, little is known about how exactly the dietary information is remotely transmitted from the gut to the neurons. To address these unsolved questions, it is critical to efficiently quantify the effects of various nutritional and genetic conditions on this nutrition-dependent hyperarborization. For this purpose, we developed DeTerm, a software program for automatic detection of dendritic branch terminals (Figures 1A and 1B; Kanaoka et al., 2019).

Here we show that class IV neurons increase their dendrite density on a low-yeast diet (LYD) compared to a high-yeast diet (HYD) due to a concurrent deficiency in vitamins, metal ions, and cholesterol. We then identified an extrinsic factor and an intracellular signaling axis that jointly enable class IV neurons to respond to the LYD nutritional status. On LYD, Akt and its upstream receptor tyrosine kinase Ror in the neuron are required for the hyperarborization. In a paracrine fashion, Wingless (Wg) produced by muscles activates Akt by way of Ror and contributes to the hyperarborization. In muscles of larvae on the HYD, Stat92E, a transcription factor in the JAK/STAT pathway, was more highly expressed and negatively regulated *wg* expression, whereas the LYD resulted in lower expression of *Stat92E*, hence higher expression of *wg*. Moreover, a ligand of the JAK-STAT pathway Unpaired 2 (Upd2) also plays a role in regulating the hyperarborization. Together, our studies illustrate how a group of nutrients in the food impact neuronal development through the Wg-Ror-Akt pathway between the neuron and closely located muscles, and that this short range-pathway is regulated by a systemic Upd2-Stat92E pathway between the fat body and muscles. As for the neuronal function, we found that LYD blunted light responsiveness of class IV neurons and larval light avoidance behavior, which may help larvae optimize their survival strategies under low-nutritional conditions.

## RESULTS

### A mixture of vitamins, metal ions and cholesterol ameliorates the hyperarborization

Our analysis using the software program called DeTerm established that both the number of branch terminals per neuron and the density of terminals (terminal number/arbor size) were higher on LYD than on HYD (Figures 1C-1E). In addition to these box plots, we drew two-dimensional plots with the dendritic area on the X-axis and the number of branch terminals on the Y-axis, and the numerical features of dendrites of class IV neurons on HYD and those on LYD were clearly separated (Figure 1F). Therefore, in the subsequent analyses, we mainly focused on the density of terminals (Figure 1E) and the separation in 2D plots (Figure 1F) to evaluate the hyperarborization phenotypes.

Yeast is one of the main ingredients in *Drosophila* laboratory foods, and it has been primarily considered as a source of amino acids. We suspected the possibility that LYD is deficient in amino acids and that is the cause of the phenotype. Therefore, we first examined whether supplementation of LYD with amino acids would ameliorate the hyperarborization. However, the addition of an essential amino acid solution, an amino acid mix, or peptone resulted in only slight or no restoration of the phenotype (Figure 1—figure supplement 1B-1L, and see details in the legend) To more comprehensively search for nutrients responsible for the hyperarborization, we used fractions of a fully chemically defined or holidic medium for *Drosophila* (Piper et al., 2014, 2017) and examined which fraction or which combinations of the fractions were able to ameliorate the phenotype (Figure 1—figure supplement 1A). Addition of four fractions other than amino acids, which comprise vitamins (V), metal ions (M), cholesterol (C), and other ingredients (nucleic acids and lipid-related metabolites: O), to LYD significantly rescued the hyperarborization (Figures 1G, 1H, 1J and 1K; see also the legend of Figure 1J). We named this diet LYD + VMCO. Further supplementation of amino acids to LYD + VMCO did not improve the degree of the rescue (Figures 1G-1K). Importantly, the phenotype was not restored without any one of three fractions, namely, vitamins, metal ions and cholesterol (Figures 1L-1R). On the other hand, the fraction designated other ingredients was dispensable for amelioration of the phenotype (Figures 1S-1U). Larval developmental timing on LYD + VMC was comparable to that on LYD (Figure 1—figure supplement 1A); nonetheless the phenotype was blunted on LYD + VMC. Therefore, these data suggest that the hyperarborization was caused by a concomitant deficiency in vitamins, metal ions, and cholesterol, rather than by extended larval development on LYD.

### Akt and receptor tyrosine kinase Ror are required in class IV neurons to hyperarborize their dendrites

To investigate the molecular mechanism underlying the hyperarborization phenotype on LYD, we focused on intracellular signaling factors that have been reported to sense nutritional status in other cellular contexts. Thus, we examined whether class IV neuron-specific knockdowns of any of these factors would affect the phenotype (Figures 2A-2J and Figure 2—figure supplement 1A-1V). We found that knockdown of *Akt kinase* (*Akt*) ameliorated the phenotype, although expression of one of the short hairpins strongly reduced dendrite length irrespective of the diets (Figures 2A-2J). In addition, we knocked down genes that constitute the signaling pathways of Akt (Figure 2—figure supplement 1W-1AT), and found that inhibition of TOR signaling components, *Target of rapamycin* (*Tor*), *Ribosomal protein S6 kinase* (*S6k*), or *Tif-IA* also ameliorated the phenotype (Figure 2—figure supplement 1W-1Y, 1AC-1AE, 1AI-1AK and 1AO-1AQ). A similar effect caused by inhibition of *Tor* was described by Poe et al (2020). On the other hand, the results of knocking down the known upstream regulators of Akt, *Insulin-like receptor* (*InR*) or *Anaplastic lymphoma kinase* (*Alk*), were difficult to interpret (Figure 2—figure supplement 1A-1F, 1AB, 1AH, 1AN and 1AT, and see details in the legend).

**Figure 2.**
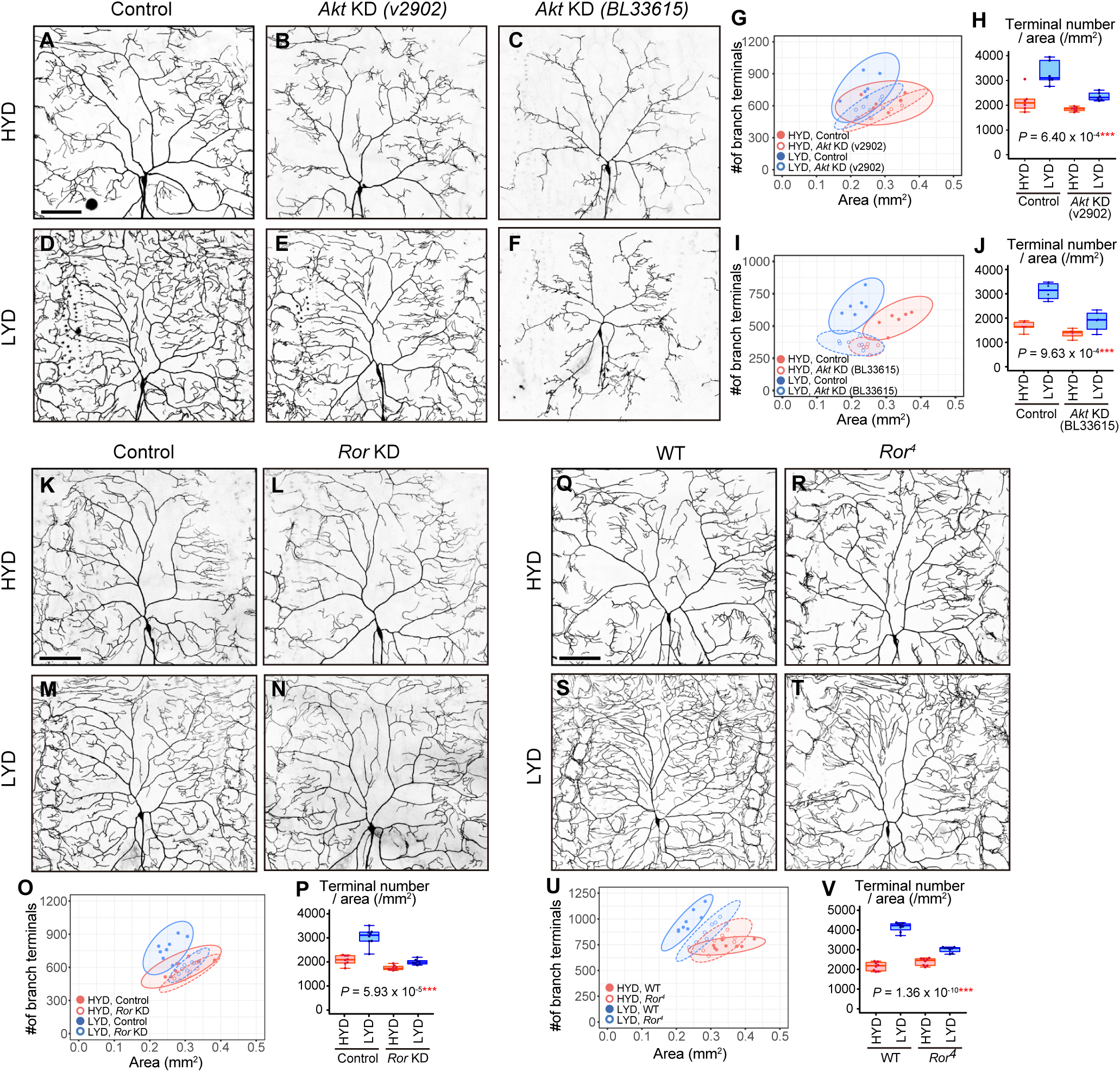
Akt and receptor tyrosine kinase Ror are required in class IV neurons to hyperarborize their dendrites. (A-F) Images of control ddaC neurons (A and D), or *Akt* knockdown ddaC neurons using *UAS-Akt RNAi^v2902^* (B and E) or *UAS-Akt RNAi^BL33615^* (C and F), on HYD (A-C) or LYD (D-F). (G-J) Quantitative analysis of effects of *Akt* knockdown using *UAS-Akt RNAi^v2902^* (G and H) or *UAS-Akt RNAi^BL33615^* (I and J). (G and I) Two-dimensional plots. (H and J) Densities of branch terminals. The differences between HYD and LYD in control and *Akt* knockdown ddaC neurons are significantly different as indicated by the P-values (Two-way ANOVA, n= 6). (K-P) Images of control (K and M) or *Ror* knocked down ddaC (L and N) on HYD (K and L) or LYD (M and N). Two-dimensional plot (O) and densities of branch terminals (P). The differences between HYD and LYD in control and *Ror* knocked down ddaC are significantly different as indicated by the P-value (Two-way ANOVA, n= 8). (Q-V) Images of ddaC in wild-type (WT; Q and S) or *Ror^4^* mutant larvae (R and T) on HYD (Q and R) or LYD (S and T). Two-dimensional plot (U) and densities of branch terminals (V). The differences between HYD and LYD in WT and *Ror^4^* mutant larvae are significantly different as indicated by the P-value (Two-way ANOVA, n= 8). Boxplots in (H, J, P, and V) are depicted as in Figure 1C. *P < 0.05, **P < 0.01, and ***P < 0.001. Scale bars, 100μm. Figure supplement for figure 2: Figure supplement 1. Contributions of intracellular signaling factors or Akt signaling components to the hyperarborization

Various secreted factors are known to function as inter-organ communication factors in response to nutritional conditions (Droujinine and Perrimon, 2016). We therefore hypothesized that, in larvae on LYD, class IV neurons receive signaling molecules from other tissues, leading to the hyperarborization via the Akt/Tor signaling pathway. As candidate receptors upstream of Akt, we focused on receptor tyrosine kinases (RTKs; Sopko and Perrimon, 2013). From knockdown screening of 20 RTK genes, we found that class IV neuron-specific knockdown of *Ror* suppressed the hyperarborization (Figures 2K-2P). We also observed class IV neurons in *Ror^4^* null mutant larvae and found that they recapitulated the result of the *Ror* knockdown (Figures 2Q-2V). These results suggest that Ror and Akt are required in class IV neurons to hyperarborize their dendrites on LYD.

### Wg in muscles is more highly expressed on LYD and promotes dendritic branching of class IV neurons

Ror binds to Wnt ligands and triggers intracellular signaling cascades (Ripp et al., 2018; van Amerongen and Nusse, 2009). We therefore knocked down *wingless* (*wg*), *Wnt2*, *Wnt4*, or *Wnt5* in either of the two tissues adjacent to class IV neurons: epidermal cells and muscles. We observed that *wg* knockdown in muscles using either *Mhc-Gal4* or *mef2-Gal4* suppressed the hyperarborization phenotype (Figures 3A-3F and Figure 3—figure supplement 1A-1F). In contrast, epidermal knockdown of *wg* had no effect on the phenotype (Figure 3—figure supplement 1G-1L). The requirement of Wg for the hyperarborization was further confirmed by the finding that the hyperarborization effect was dampened in class IV neurons in the whole-body *wg* mutant (hypomorphic *wg^1^*/amorphic *wg ^l-8^*; Figures 3G-3L).

**Figure 3.**
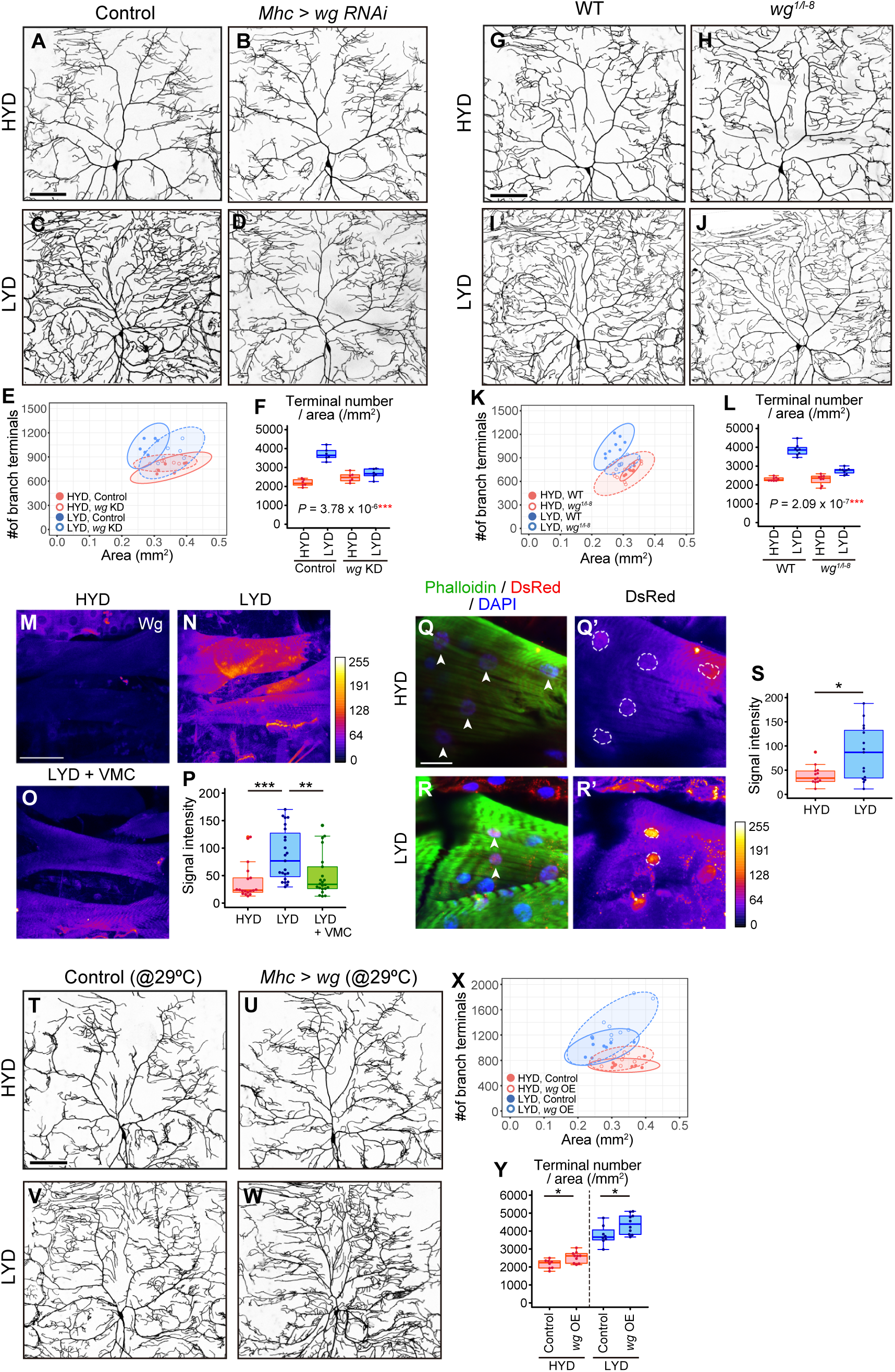
Wg in muscles is expressed more highly on LYD and promotes dendritic branching of class IV neurons. (A-F) Images of ddaC neurons in control larvae (A and C) or larvae with *wg* knockdown in muscles (B and D), on HYD or LYD. Two-dimensional plot (E) and densities of branch terminals (F). The differences between HYD and LYD in control larvae and larvae with *wg* knockdown in muscles are significantly different as indicated by the P-value (Two-way ANOVA, n= 6). (G-L) Images of ddaC neurons in WT (G and I) or *wg^1/l-8^* larvae (H and J) on HYD or LYD. Two-dimensional plot (K) and densities of branch terminals (L). The differences between HYD and LYD in WT and *wg^1/l-8^* larvae are significantly different as indicated by the P-value (Two-way ANOVA, n = 8). (M-P) Muscles in larvae reared on HYD (M), LYD (N), or LYD + VMC (O) were stained for Wg. The signal intensities are represented by the indicated color code. (P) Quantification of the mean Wg immunofluorescence intensity in muscle 9, one of the closest muscles to the ddaC neuron (Steel Dwass test, n = 18-23). (Q-S) Images of *wg-Gal4* muscles driving the expression of RedStinger, which is DsRed tagged with a nuclear localization signal. Muscles, on HYD (Q) or LYD (R) were stained with phalloidin (green), an antibody to DsRed (red), and DAPI (blue). The signal intensities of DsRed are represented by the indicated color code, and white dashed circles indicate outlines of nuclei (Q’and R’). (S) Quantification of DsRed intensity in nuclei of muscle 9 (Wilcoxon-Mann-Whitney test, n = 13-15). (T-Y) Images of ddaC neurons in control larvae (T and V) or larvae with *wg* overexpression in muscles (U and W) on HYD or LYD. Two-dimensional plot (X) and densities of branch terminals (Y; Wilcoxon-Mann-Whitney test, n = 8-10). Boxplots in (F, L, P, S, and Y) are depicted as in Figure 1C. *P < 0.05, **P < 0.01, and ***P < 0.001. Scale bars, 100μm (A-D, G-J, M-O, and T-W), 25μm (Q-R’). Figure supplement for figure 3: Figure supplement 1. Wg from muscles, but not from epidermal cells, contributes to the hyperarboriztion phenotype

We then examined whether Wg is differentially expressed in muscles between larvae reared on HYD and those on LYD. Immunostaining using an anti-Wg antibody showed stronger signals in LYD-fed larvae (Figures 3M, 3N, and 3P). These stronger signals indeed represented increased amounts of endogenous Wg because knocking down *wg* decreased the intensity (Figure 3—figure supplement 1M-1Q). We also asked whether *wg* expression is up-regulated on LYD at the transcriptional level. We expressed RedStinger, DsRed tagged with a nuclear localization signal, under the knocked-in *wg-Gal4* driver that reflects the endogenous expression pattern of *wg* (Bosch et al., 2020). Nuclear RedStinger signals in muscles were stronger in larvae on LYD (Figures 3Q-3S), indicating that LYD up-regulated *wg* transcription compared to HYD. We further tested whether muscle-derived Wg promotes dendritic branching of class IV neurons. For this purpose, we overexpressed *wg* in muscles and found that those larvae increased the number of dendritic terminals per neuron on both HYD and LYD (Figures 3T-3Y), strengthening the role of the muscle–class IV neuron communication in hyperarborization. Importantly, addition of vitamins, metal ions, and cholesterol to LYD significantly suppressed the up-regulation of Wg on LYD (Figures 3M-3P). Together with the effect of these compounds on dendritic branching (Figures 1S-1U), we hypothesized that *wg* expression in muscles is enhanced by a concurrent deficiency in vitamins, metal ions, and cholesterol in LYD, and that muscle-derived Wg promotes dendritic branching of class IV neurons.

### Wg-Ror-mediated activation of Akt in class IV neurons evokes the hyperarborization

Wnt signaling is engaged in diverse contexts of neuronal development and regeneration (Green et al., 2014; He et al., 2018; Endo and Minami, 2018; Nye et al., 2020; Weiner et al., 2020). In previous study in *Drosophila*, responses to dendrite injuries were investigated using class I and IV neurons. This showed that Ror, a seven-pass transmembrane receptor Frizzled (Fz), and downstream components including Disheveled (Dsh) and Axin (Axn) are required for dendrite regeneration (Nye et al., 2020). We therefore examined whether these genes and other components of Wnt signaling affect the hyperarborization phenotype (Figure 4—figure supplement 1 and Figure 4—figure supplement 2). Knocking down *fz2* significantly ameliorated the hyperarborization (Figure 4—figure supplement 1A-1J). Not only *fz2* knockdown neurons, but also *fz2* null mutant neurons showed less prominent hyperarborization compared to the control neurons (Figure 4—figure supplement 1K-1N). These results are consistent with the proposed function of Ror as a Wnt co-receptor with Fz2 (Ripp et al., 2018). Among the components downstream of the receptors, knocking down *dsh* or blocking the function of *basket* (*bsk*) encoding JNK reduced the differences in the densities of dendritic terminals between the diets (Figure 4—figure supplement 2A-2C, 2H-2J, 2O, 2P, 2U, and 2V). However, these results need careful interpretation (see the Figure 4—figure supplement 2 legend). Altogether, our results suggest that among the known components of Wnt signaling in *Drosophila*, Fz2 cooperates with Ror in transducing the external signal to evoke the hyperarborization.

Ror is also reported to activate the PI3K/Akt/mTor signaling pathway in lung adenocarcinoma or multiple myeloma (Liu et al., 2015; Frenquelli et al., 2020). We therefore hypothesized that Wg-Ror signaling activates Akt signaling in class IV neurons on LYD, leading to the hyperarborization. To test this hypothesis and to clarify the relationship between Wg-Ror and Akt at the molecular level, we examined how genetic manipulations of Wg-Ror signaling affect p-Akt levels in class IV neurons (Figures 4A-4J). We first examined how the p-Akt level in class IV neurons differs between larvae reared on HYD and LYD. Immunostaining showed that the p-Akt level in class IV neurons was higher on LYD than on HYD (Figures 4A, 4A’, 4C, 4C’ and 4E). In contrast, *Ror* knockdown neurons from larvae on LYD showed reduced p-Akt levels compared to those on HYD (Figures 4B, 4B’, 4D, 4D’ and 4E). Furthermore, *wg* overexpression in muscles increased the p-Akt level in class IV neurons on HYD (compare Figure 4F’ with 4G’), which became comparable to the level on LYD (compare Figure 4G’ with 4I’; see quantification in Figure 4J). These results suggest that Akt signaling in class IV neurons is activated by muscle-derived Wg, and this activation is mediated by Ror in the neurons.

**Figure 4.**
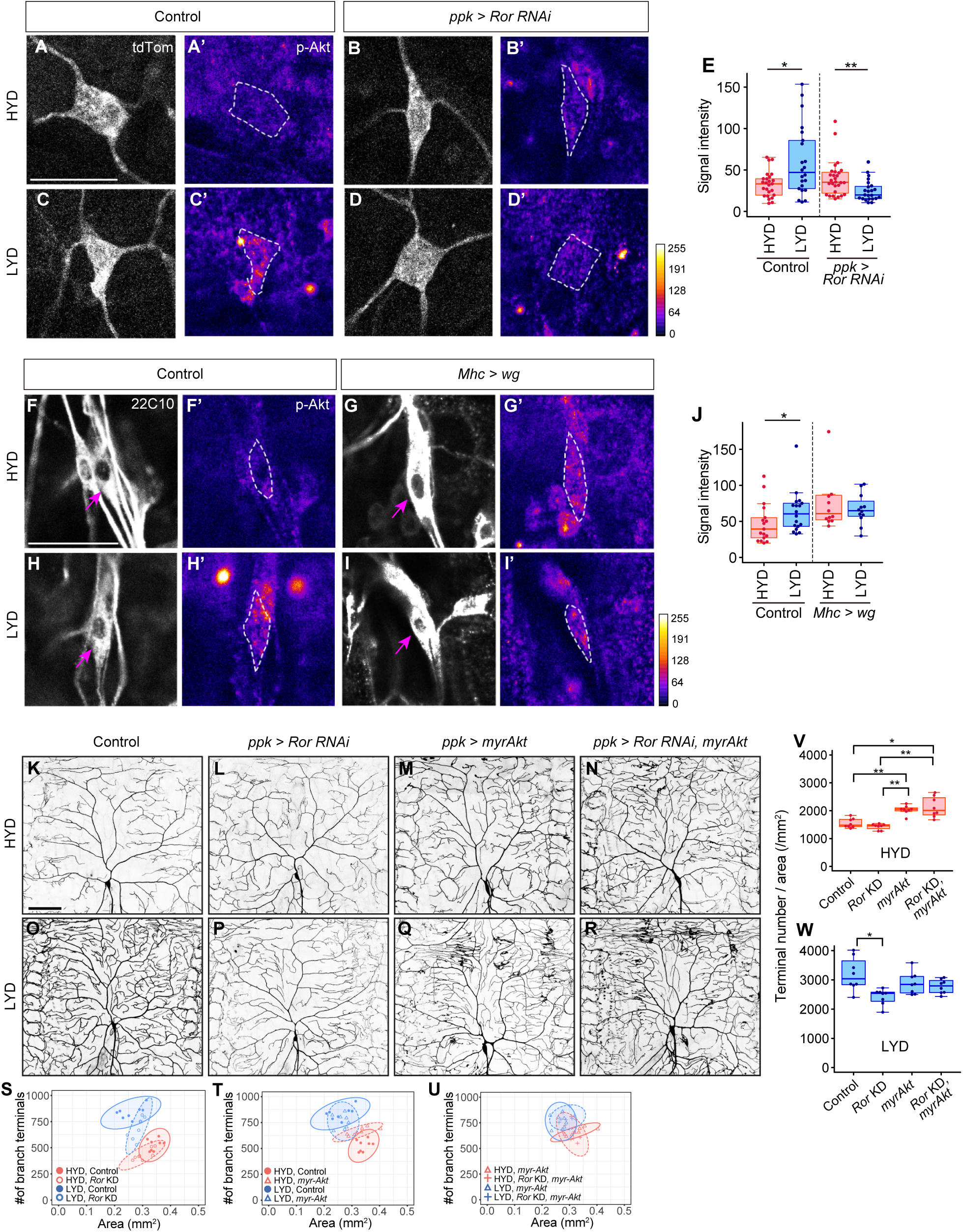
Wg-Ror-mediated activation of Akt in class IV neurons evokes the hyperarborization. (A-E) Control ddaC (A, A’, C and C’) or *Ror* knockdown ddaC neurons (B, B’, D and D’) were stained for p-Akt (A’-D’) and co-imaged with a class IV marker *ppk-CD4:tdTom* (A-D). The signal intensities of p-Akt correspond to the indicated color code, at right, and white dashed circles indicate the cell bodies of ddaC neurons. (E) Quantification of p-Akt intensity in cell bodies of control or *Ror* knocked down ddaC neurons (Wilcoxon-Mann-Whitney test, n = 22-28). (F-J) ddaC in control larvae (F, F’, H and H’) or larvae with *wg* overexpression in muscles (G, G’, I and I’) were stained for a pan-sensory neuron marker (22C10; F-I) and for p-Akt (F’-I’). (F-I) Magenta arrows indicate the cell bodies of ddaC neurons. (F’-I’) The intensities of p-Akt signals correspond to the indicated color code, at right, and white dashed circles indicate the cell bodies of ddaC neurons. (J) Quantification of p-Akt intensity in control larvae or larvae with *wg* overexpression in muscles (Wilcoxon-Mann-Whitney test, n = 12-18). (K-U) Images of control (K and O), *Ror* knockdown ddaC neurons (L and P), ddaC neurons with expression of *myrAkt,* which is a constitutive active form of *Akt* (M and Q), or *Ror* knockdown ddaC neurons with *myrAkt* expression (N and R) on HYD or LYD. (S-W) Quantitative analysis of combinatorial effects of *Ror* knockdown and *myrAkt* expression. (S-U) Two-dimensional plots. (V and W) Densities of branch terminals on HYD (V) or LYD (W) (Steel Dwass test, n = 8-9). Boxplots in (E, J, V and W) are depicted as in Figure 1C. *P < 0.05 and **P < 0.01. Scale bars, 25μm (A-D’ and F-I’), 100μm (K-R). Figure supplements for figure 4: Figure supplement 1. Fz2, a receptor for Wnt proteins, is required in class IV neurons to hyperarborize their dendrites Figure supplement 2. Effects of inhibiting intracellular Wnt signaling components on hyperarborization

We further examined whether the activation of Akt itself evokes hyperarborization even in the absence of the upstream Ror-mediated signaling (Figures 4K-4W). Expression of myr-Akt, a constitutively activated membrane-anchored form of Akt (Stocker et al., 2002), in class IV neurons increased the terminal density even on HYD, regardless of whether *Ror* was knocked down or not (compare Figure 4K with 4M and 4N; see also 4T, 4U and 4V). This result suggests that Akt activation in the neurons plays a pivotal role for the hyperarborization. Our result is consistent with a previous finding that overexpression of the wild-type form of *Akt* causes a significant increase in dendrite coverage of the epidermis (Parrish et al., 2009).

### Downregulation of Wg expression by Stat92E suppresses hyperarborization on HYD

Given that Wg expression in muscles is higher on LYD (Figures 3M-3P), and the differential expression impacts the dendritic branching of class IV neurons (Figures 3T-3Y), we then asked how Wg expression in muscles is regulated in the nutrient-dependent manner. To search for upstream regulators of the Wg expression, we performed RNA-seq analysis on mature whole larvae that were reared on either diet. We identified differentially expressed 3854 genes between the diets (Figures 5A, Figure 5—figure supplement 1, and Supplementary File 2). Among these, we focused on a transcriptional factor in JAK/STAT pathway, *Stat92E*, which is more highly expressed on HYD than LYD (Figure 5B). Also informing our decision, it was reported that Stat92E is a negative regulator of *wg* expression in the eye imaginal disc (Ekas et al., 2006). We used a Stat92E reporter strain (Bach et al., 2007) and found that Stat92E reporter expression in muscle was higher on HYD (Figures 5C-5E). We therefore hypothesized that, in muscles of larvae on HYD, higher expression of Stat92E downregulates Wg expression, thereby suppressing the hyperarborization phenotype. To test this hypothesis, we knocked down *Stat92E* in muscles, and this led to increased Wg levels compared to the control muscles on HYD (Figures 5F-5J; compare 5F with 5G). Furthermore, knocking down *Stat92E* or *hopscotch* (*hop*) encoding JAK in muscles promoted hyperarborization of class IV neurons in larvae reared on HYD (Figures 5K-5T and Figure 5—figure supplement 2A, 2B, 2F, 2G, 2K and 2O). In contrast, overexpression of *hop* in muscles ameliorated the hyperarborization on LYD (Figures 5U-5Z). These results indicate that JAK/STAT signaling contributes to the downregulation of *wg* expression in muscles and the suppression of dendritic hyperarborization on HYD (Figure 7).

**Figure 5.**
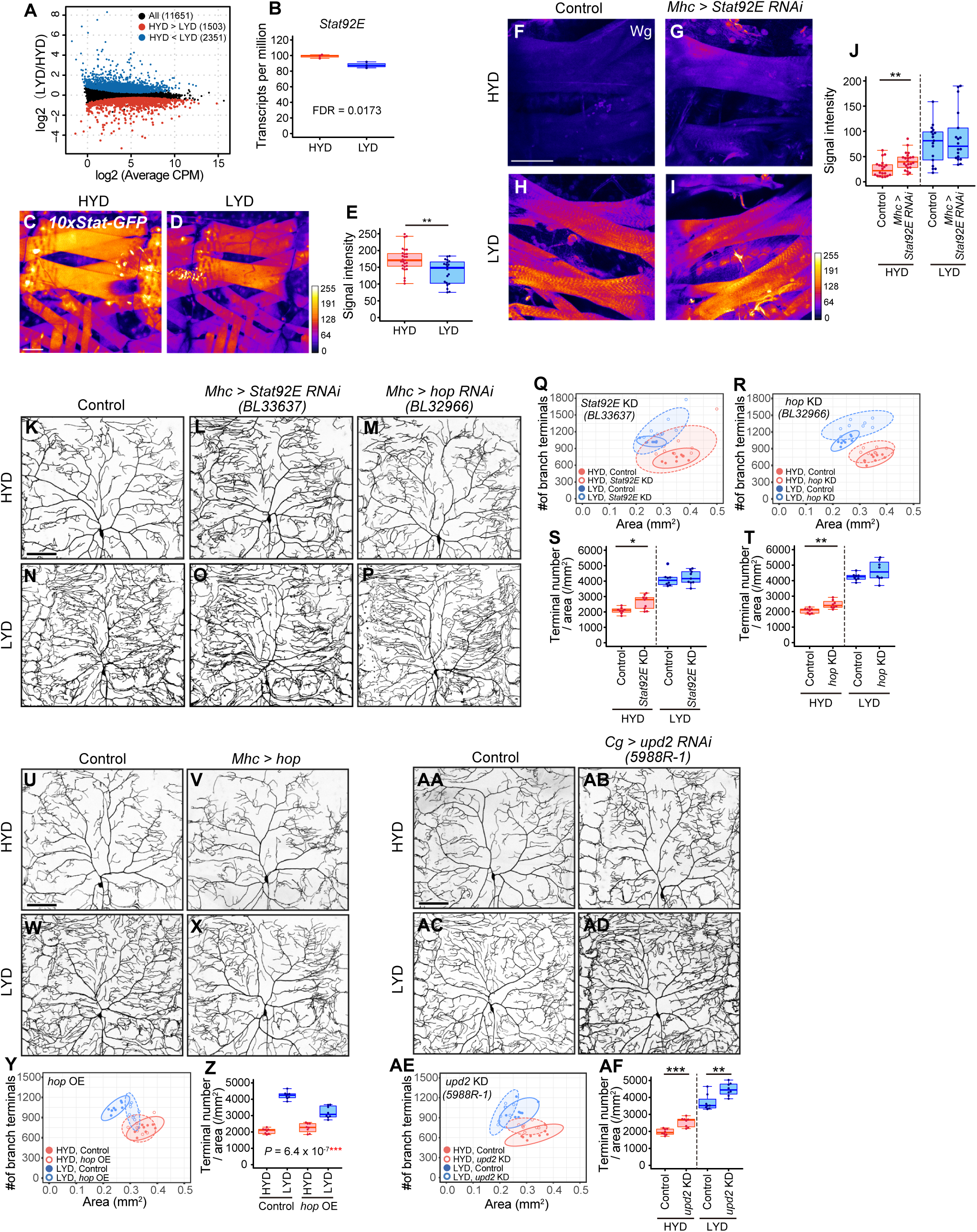
Downregulation of Wg expression by Stat92E on HYD suppresses the hyperarborization phenotype. (A and B) Plots of whole-body RNA-seq of wandering 3^rd^ instar larvae reared on HYD or LYD. (A) The fold change (LYD/HYD) in read counts was plotted against average counts per million mapped reads (CPM) for HYD and LYD. Dots that are statistically supported (FDR ≤0.05) are colored (red for HYD>LYD and blue for HYD<LYD). (B) Plot of transcripts per million (TPM) of *Stat92E*. Adjusted P-value with Benjamini & Hochberg correction (FDR) is indicated. (C-E) Muscles of *10x Stat-GFP* larvae on HYD (C) or LYD (D). The signal intensities of GFP correspond to the indicated color code. (E) Quantification of 10x Stat-GFP intensity in muscle 9 (Student’s t-test, n = 20-27). (F-J) Muscles of control larvae (F and H) or larvae with *Stat92E* knockdown in muscles (G and I) on HYD or LYD were stained for Wg. The signal intensities correspond to the indicated color code. (J) Quantification of the mean Wg immunofluorescence in muscle 9 (Wilcoxon-Mann-Whitney test, n = 17-22). (K-T) Images of ddaC neurons in control larvae (K and N), larvae with *Stat92E* knockdown in muscles (L and O), or larvae with *hop* knockdown in muscles (M and P), on HYD or LYD. Two-dimensional plots (Q and R) and densities of branch terminals (S and T; Wilcoxon-Mann-Whitney test, n = 8-9). (U-Z) Images of ddaC neurons in control larvae (U and W) or larvae with *hop* overexpression in muscles (V and X), on HYD or LYD. Two-dimensional plot (Y) and densities of branch terminals (Z). The differences between HYD and LYD in control larvae and larvae with *hop* overexpression in muscles are significantly different as indicated by the P-value (Two-way ANOVA, n= 8). (AA-AF) Images of ddaC neurons in control larvae (AA and AC) or larvae with *upd2* knockdown in the fat body and hemocytes (AB and AD), raised on HYD or LYD. Two-dimensional plot (AE) and densities of branch terminals (AF; Wilcoxon-Mann-Whitney test, n = 8). Control data in (R) and (T) are shared with (Y) and (Z). Boxplots in (B, E, J, S, T, Z and AF) are depicted as in Figure 1C. *P < 0.05, **P < 0.01, and ***P < 0.001. Scale bars, 100μm. Figure supplements for figure 5: Figure supplement 1. Enriched terms in functional annotation clustering of differentially expressed genes depending on diets in whole body RNA-seq data Figure supplement 2. Effects of inhibiting components of JAK/STAT pathway on hyperarborization

It was previously reported that Upd2 secreted from the fat body activates JAK/STAT signaling through transmembrane receptor Domeless (Dome) in GABAergic neurons, which project onto insulin producing cells (IPCs), thereby regulating larval growth in a nutritional-status-dependent manner (Rajan and Perrimon, 2012). It has also been shown that the secretion of Upd2 or Upd3 from hemocytes promotes the expression of a *Stat92E* reporter in muscle (Yang et al., 2015). These studies prompted us to address whether any Upds from the fat body or hemocytes, and Dome in muscles, contribute to the hyperarborization phenotype. Knocking down *dome* in muscles hardly affected the hyperarborization (Figure 5—figure supplement 2C-2E, 2H-2J, 2L-2N and 2P-2R), whereas knocking down *upd2,* but not *upd* or *upd3,* in the fat body and hemocytes resulted in an increased terminal density on HYD (Figures 5AA-5AF and Figure 5—figure supplement 2S-2AN). This effect of *upd2* knockdown in the fat body and hemocytes is similar to that of *hop* or *Stat92E* knockdown (Figures 5K-5T) and that of *wg* overexpression in muscles (Figures 3T-3Y). These results are suggestive of the role of fat body (and hemocytes)–muscle inter-organ communication through a Upd2-Stat92E pathway in suppressing the hyperarborization phenotype on HYD (Figure 7).

### LYD blunts light responsiveness of class IV neurons and larval light avoidance behavior

Class IV neurons sense noxious thermal, mechanical, and light stimuli (Chin and Tracey, 2017). We therefore examined how our dietary conditions affect the electrophysiological activity of class IV neurons and larval behavior (Figure 6). First, we compared firing activities of class IV neurons in larvae that were reared on either HYD or LYD. As a noxious stimulus, we illuminated entire arbors of recorded neurons with blue light (Xiang et al., 2010; Terada et al., 2016). We used extracellular recording to monitor both spontaneous and evoked activities (Figures 6A-6C). The frequency of spontaneous firing was higher in class IV neurons from larvae reared on LYD than on HYD (Figure 6D). Regarding the response to the light stimulus, all relevant parameters, i.e., the firing frequency, the change amount, and the change rate, were lower on LYD than on HYD (Figures 6E-6G, see definition of the parameters in the legend), indicating that class IV neurons on LYD are less sensitive to the stimulus. Next, we examined whether the blunted light responsiveness of the neurons affects larval avoidance behavior. It was reported that *Drosophila* larvae prefer dark place to avoid noxious light and this light avoidance behavior requires the activity of class IV neurons (Yamanaka et al., 2013; Imambocus et al., 2022). We speculated that the blunted light responsiveness of larvae on LYD may cause declines in their light avoidance behavior, and this may allow larvae to continue their search for high-nutrient food. To address this possibility, we conducted light/dark choice assays in which larvae reared on HYD or LYD were allowed to choose between dark and bright areas. We found that both foraging 3^rd^ instar and wandering 3^rd^ instar larvae on LYD showed lower preference for dark places than larvae on HYD (Figures 6H and 6I). Furthermore, when *Ror* was knockdown in class IV neurons, differences in light avoidance behavior between the diets tended to be smaller than the control larvae (Figures 6H and 6I). Our results suggest that the hyperarborization of class IV neurons is associated with blunted light avoidance behavior.

**Figure 6.**
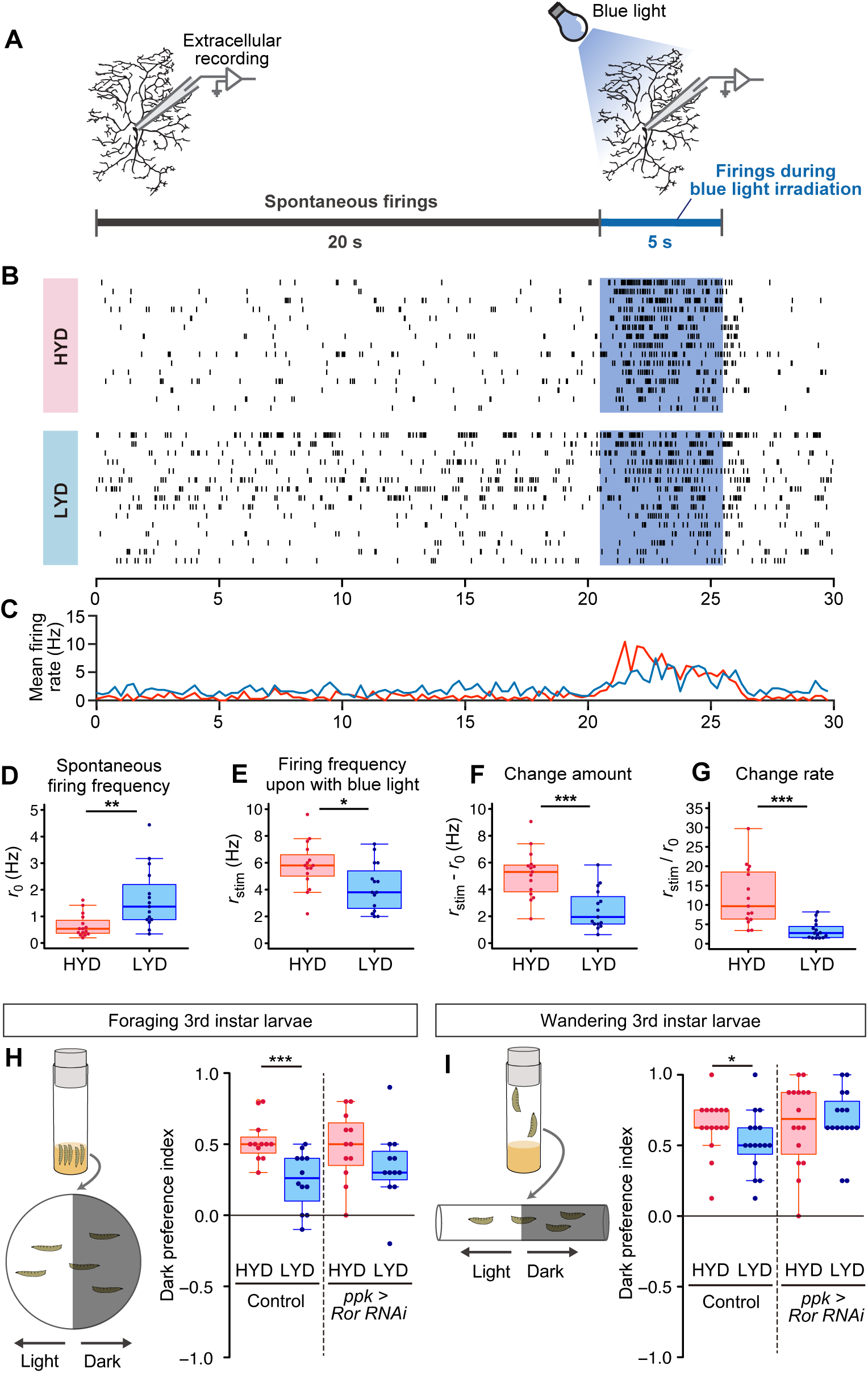
LYD blunts light responsiveness of class IV neurons and larval light avoidance behavior. (A) A schematic diagram outlining the electrophysiological analysis. Firing activities of class IV neurons v’ada were recorded by measuring the extracellular membrane potential. After spontaneous firings were recorded for about 20 s, activities during blue light irradiation were monitored for 5 s. (B and C) Firing activities of class IV neurons on HYD or LYD. (B) Raster plots of firing. Blue shading indicates the 5 s blue light irradiation. Each row in the plots represents the data for a single cell. (C) Peristimulus time histograms calculated at 250-ms bins, on HYD (red line) or LYD (blue line). (D-G) Quantitative analysis of the firing activities. (D) Spontaneous firing frequency. (E) Firing frequency during blue light irradiation. (F) Change amount of the firing response to the blue light stimulus calculated by subtracting [spontaneous firing frequency] from [firing frequency during blue light irradiation] (G) Change rate of the firing response to the blue light stimulus calculated by dividing [firing frequency during blue light irradiation] by [spontaneous firing frequency]. (Wilcoxon-Mann-Whitney test, n = 15) (H and I) Schematic diagram of light/dark choice assays and dark preference index of foraging 3^rd^ instar larvae on agar plates (H), and wandering 3^rd^ instar larvae in plastic tubes (I). Control larvae and larvae with *Ror* knockdown in class IV neurons were tested. (Wilcoxon-Mann-Whitney test, n = 12-16) Boxplots in (D-I) are depicted as in Figure 1C. *P < 0.05, **P < 0.01, and ***P < 0.001.

## DISCUSSION

Collectively, our studies illustrate how selective nutrients in the food impact neuronal development through inter-organ signaling (Figure 7). Yeast has long been considered as a rich source of amino acids for *Drosophila*; however, our results suggest that class IV neurons increase their dendrite density on the LYD due to a combined deficiency in vitamins, metal ions, and cholesterol. This result is unexpected, because previous studies on nutrition-dependent cell growth has focused primarily on the TOR signaling pathway, which is activated by amino acids (Gonzalez et al., 2020; Liu and Sabatini, 2020). The addition of the above nutrient trio to LYD did not restore the dendrite density to the same level as HYD. This may indicate that the balance of concentrations among these nutrients was not optimized or that unknown nutrients may need to be added along with these nutrients. In addition, we showed that up-regulation of Wg expression in muscle on LYD was suppressed by supplementation of these components to the diet. This regulation may be achieved at least at a transcriptional level possibly by an interplay between a hypothetical nutrient-responsive module in the cis-element of *wg* and transcription factors and/or epigenetic machineries that require vitamins, vitamin-derived metabolites, metal ions and cholesterol (Harris et al. 2016; Kambe et al., 2016). Further investigations are necessary to understand the detailed molecular mechanisms underlying the combined effects of these components on *wg* expression. It has been reported that increasing or decreasing the amount of yeast in foods causes various responses in *Drosophila* (Bass et al, 2007; Okamoto and Nishimura, 2015), and the approach used in this study may help to clarify which nutrients are the key factors that cause those responses.

**Figure 7.**
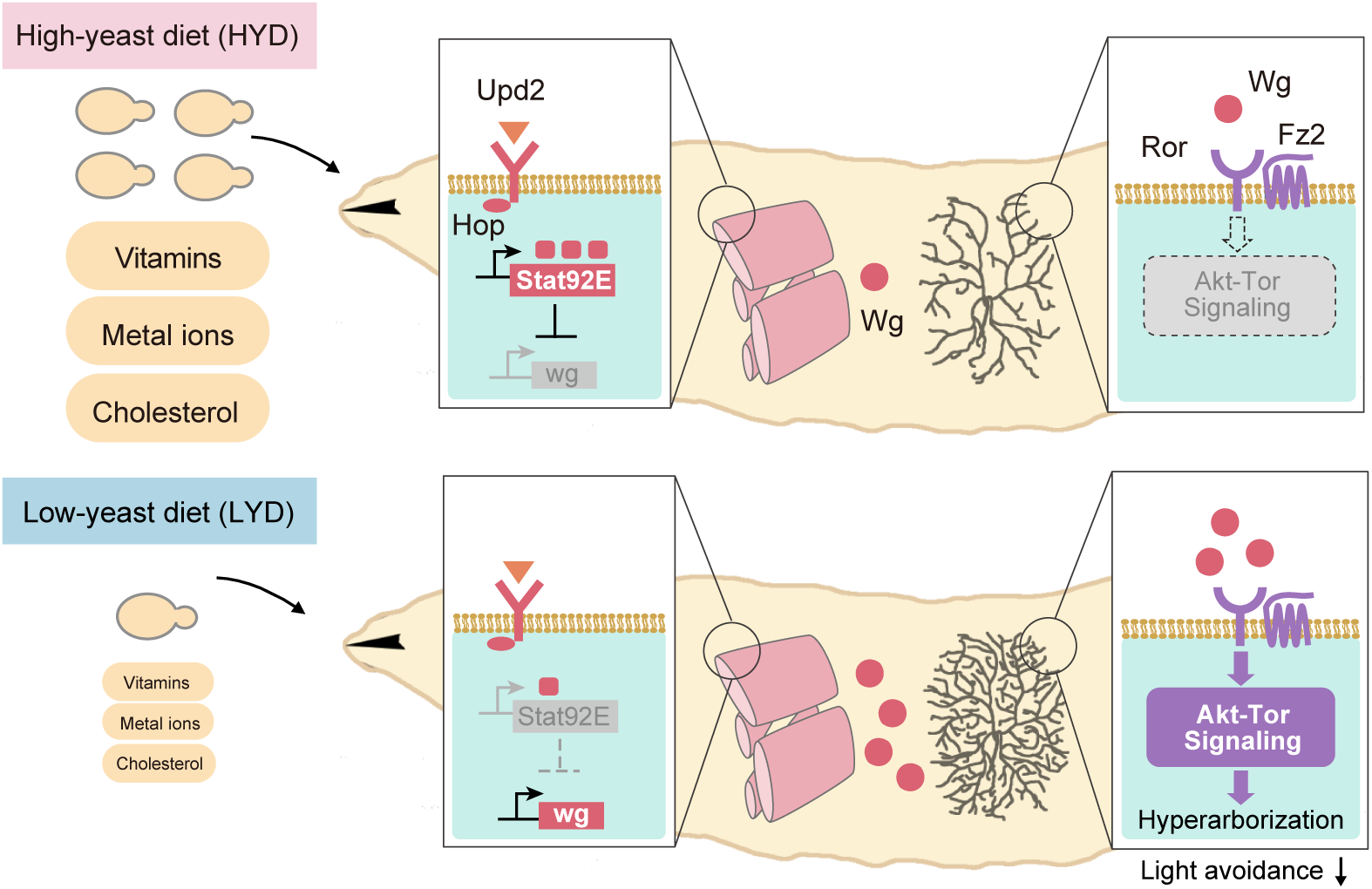
Model of the low-nutrient dependent dendritic hyperarborization. See results and discussion for details.

### Inter-organ Wg/Ror/Akt signaling-mediated the hyperarborization phenotype

Previous studies demonstrated the coordinate growth control of dendrites of class IV neurons and the epidermis (Parrish et al., 2009; Jiang et al., 2014), from which a separate model underlying dendritic hyperarborization has evolved (Poe et al., 2020). In that model, stress sensor FoxO is expressed less in neurons than in neighboring epidermal cells, which results in lower levels of autophagy and less suppression of Tor signaling in the neuron, thereby ensuring dendritic growth even under a low-yeast condition (Poe et al., 2020). In contrast to the above model, our model highlights the signaling between neurons and another adjacent tissue: muscles secrete Wg, while class IV neurons express the receptor complex Ror-Fz2 on their cell surface. Therefore, it is likely that both the extrinsic Wg-dependent mechanism and the FoxO-dependent intrinsic sparing mechanism work together to generate the hyperarborization phenotype.

Ror is mainly expressed in the nervous system. No significant abnormalities in neuronal morphology including that of the class IV neurons were observed in *Ror* mutants under standard dietary conditions (Ripp et al., 2018; Nye et al., 2020). Consistent with these reports, there was no significant difference in morphological features of dendritic arbors of class IV neurons between Ror mutant or knockdown larvae and control larvae under the nutrient-rich HYD condition (Figures 2L and 2R). Because Ror is required for the hyperarborization under the hypotrophic condition (LYD in this study; Figures 2N and 2T) and for dendrite regeneration (Nye et al., 2020), it could be an adaptive agent that copes with environmental stress or damage. A number of other RTKs, such as insulin/IGF receptors and EGFR, activate Akt in other cellular contexts such as growth and proliferation of various stem cells and mammalian cancer cells (Shim et al., 2013; Butti et al., 2018). It is likely that InR also functions upstream of Akt in class IV neurons (Parrish et al., 2009; Shimono et al., 2014; Poe et al., 2020). Future studies will explore how these various inputs are integrated by Akt to realize the nutritional status-dependent dendrite branching of class IV neurons.

The muscle is not only an energy-consuming organ, but it also plays an important role in regulating metabolic signaling through inter-organ communication with other tissues such as the brain and the fat body (Bretscher and O’Conner, 2020). In the adult stage, for example, muscle-derived Wg regulates lipid storage in the fat body (Lee et al., 2014). Our study revealed that muscle-derived Wg, which is up-regulated in response to low levels of vitamins, metal ions, and cholesterol, regulates dendrite branching of class IV neurons in the larval stage. Therefore, the muscle functions as a mediator of the nutritional status to other peripheral tissues in both growing and adult stages, and Wnt signaling may play a pivotal role in fulfilling this metabolic response function throughout the life cycle.

In our search for regulatory mechanisms of *wg* expression, we found that Stat92E reporter expression was higher in muscles on HYD than on LYD. This finding is reminiscent of Stat92E reporter expression in a population of GABAergic neurons in the adult brain, which project onto the Insulin producing cells (IPCs) and inhibit the release of Dilps (Rajan and Perrimon, 2012). This reporter expression in the GABAergic neurons also varies in a nutritional-status-dependent manner: the expression is higher on a standard laboratory food containing yeast compared to a sucrose-only condition. To address whether the key nutrients (vitamins, metal ions, and cholesterol) increase *Stat92E* reporter expression in muscles, we examined the reporter expression on LYD supplemented with or without those nutrients. The addition of all of those nutrients together to LYD did not increase the level of *Stat92E* reporter intensity (data not shown). This result contrasts with the decreased level of Wg in response to the key nutrients (Figures 3M-3P). Further investigation is required to elucidate how Wg expression in muscles is controlled at the molecular level in such a key nutrient(s)-dependent manner and how the Upd2-Stat92E pathway contributes to the entire mechanism of inter-organ communication.

### Physiological roles of class IV neurons and the hyperarborization phenotype

What are the implications of the inter-organ signaling mechanism controlling dendritic branches in the context of nutritional adaptation? It has been reported that a wide range of animals tend to take more risks when they are hungry (Symmonds et al, 2010; Filosa et al., 2016; Padilla et al., 2016; Bräcker et al., 2013). Our electrophysiological analysis indicates that class IV neurons on LYD decrease light-evoked response (Figures 6A-6G). Consistently, larvae reared on LYD displayed decreases in their dark preference, compared to those reared on HYD, and they explored bright places, which is potentially risky for their survival (Control of Figures 6H and 6I). This difference between the diets tended to become smaller once the inter-organ signaling mechanism was suppressed in class IV neurons (*ppk*>*Ror RNAi* in Figures 6H and 6I). These results imply that the hyperarborization of class IV neurons on LYD might contribute to blunting light avoidance behavior, although we cannot exclude the possibility that *Ror* knockdown might affect neuronal functions through other mechanisms than the dendritic morphological change. Our study raises the possibility that nutrient-dependent development of somatosensory neurons plays a role in optimizing a trade-off between searching for high-nutrient foods and escaping from noxious environmental threats. Although a recent study described the circuitry required for the larval light avoidance behavior, it remains unclear whether the possible modifications of neural circuits downstream of class IV neurons take part in this behavioral transition (Imambocus et al., 2022). The identification of the downstream circuits would allow further study of the relationship between nutrient-dependent neural differentiations and evoking risk-taking behavior.

In contrast to our results, a previous study reported that larvae with hyperarborized class IV neurons react more quickly to noxious heat (Poe et al., 2020). While light-induced Ca^2+^ activity in class IV neurons decreases, thermal nociceptive behavior increases during 2^nd^ and 3^rd^ instar larval periods (Jaszczak et al., 2022), which indicates that these nociceptive responses are regulated in the opposite direction or in distinct fashions. Therefore, seemingly contradictory results between the previous study and ours may be due to different regulatory mechanisms of the sensory modalities.

The relationship between nutritional status and neural development has often been studied epidemiologically (Prado and Dewey, 2014; Bhutta et al., 2017). Our study, which presents a mechanism by which quantitative changes in specific nutrients act on neuronal morphology and operate through inter-organ signaling, provides a stepping stone for future explorations of molecular mechanisms linking nutrition and development of other neuronal cell types and in other animal species.

## MATERIALS AND METHODS

### *Drosophila* strains and fly culture

Fly strains used in this study are listed in Table 1. Our stocks are usually reared on a laboratory standard diet (Watanabe et al., 2017). Adult males and virgin females that had developed on the standard diet were collected and crossed on the standard diet for 3-5 days. Then, the adults were transferred into vials containing HYD or LYD, which were identical to the semidefined medium (SDM)-based diet (8% Y) and the SDM-based diet (0.8% Y), respectively (see Supplementary File 1 and its legend in Watanabe et al., 2017). After an egg-laying interval, the adult flies were cleared in every experiment and wandering 3rd instar larvae that came out of individual diets were used. Larvae were reared under noncrowded conditions at 25 °C in all the experiments except the *wg* overexpression experiments at 29 °C. Our recombinant DNA experiments follow Kyoto University Regulations for Safety Management in Recombinant DNA Experiments under protocol # 210059.

**Table 1.**
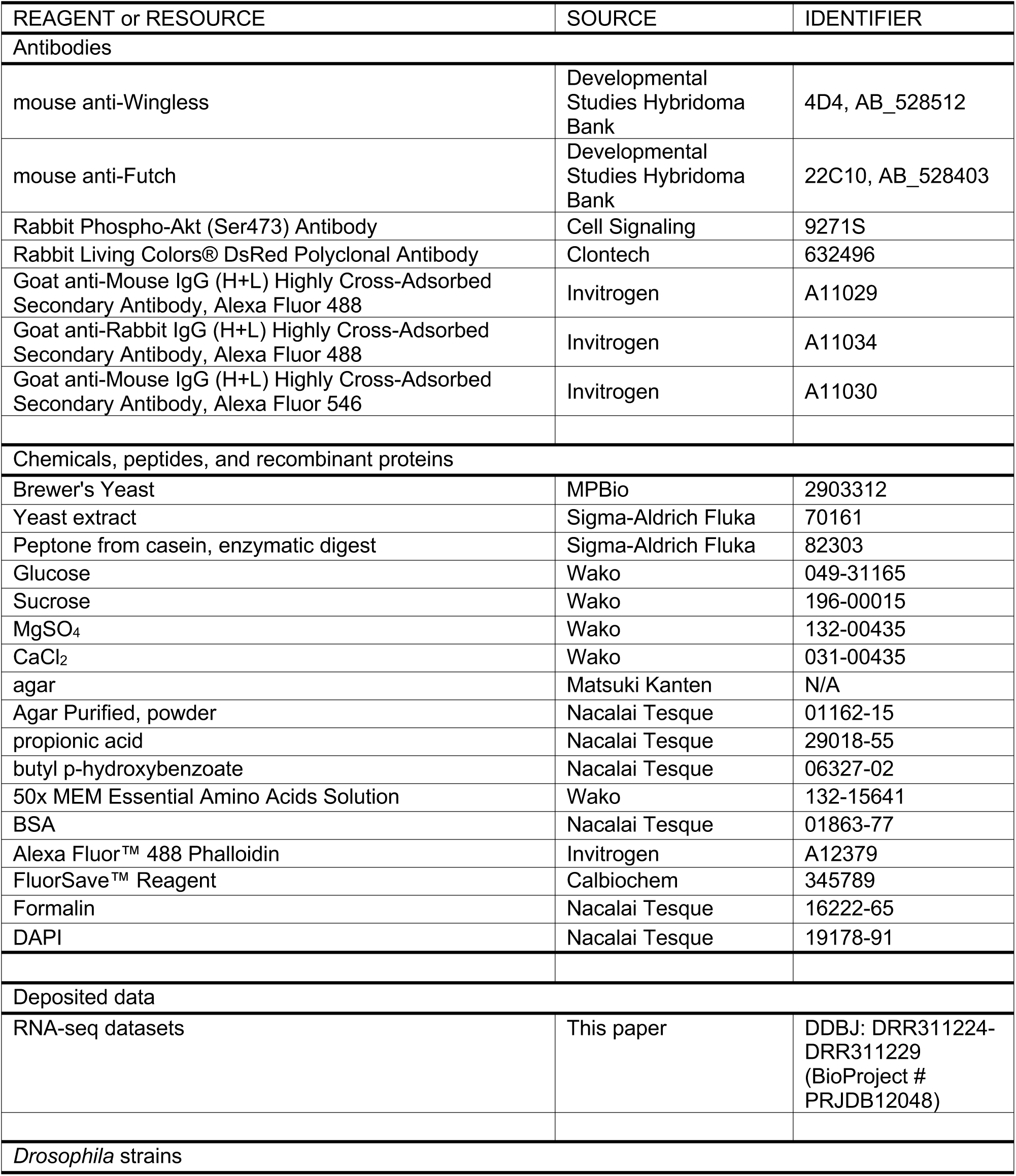
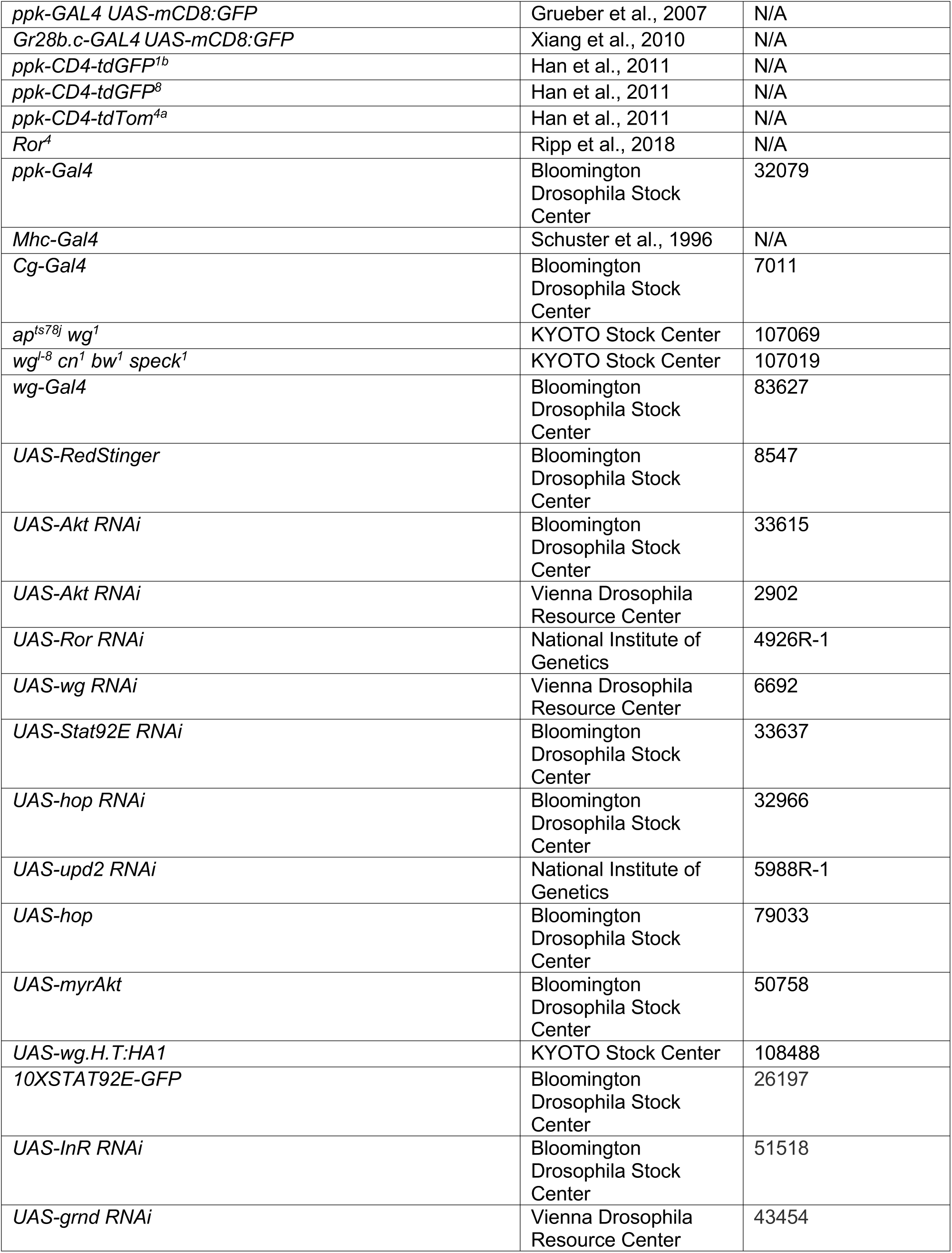
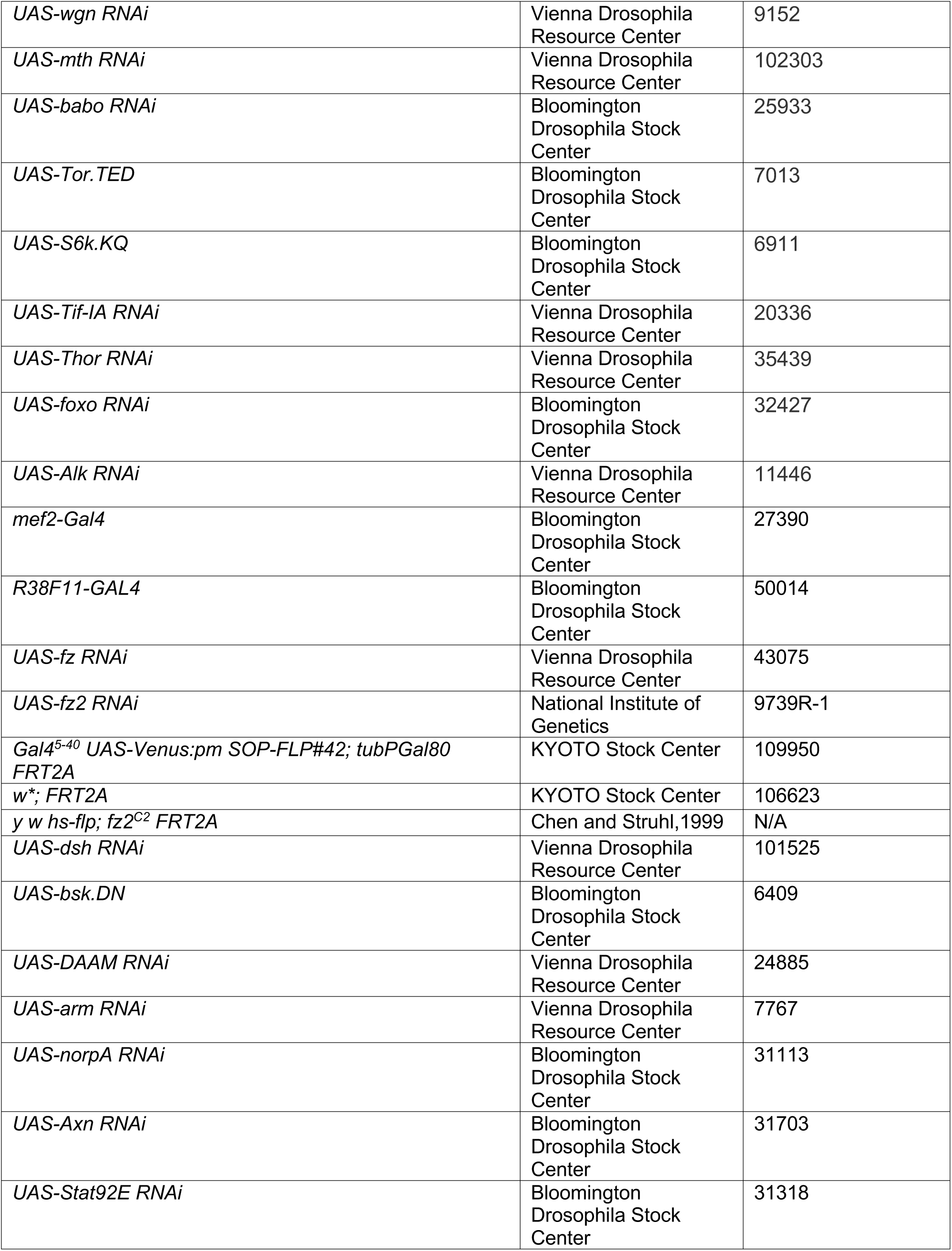
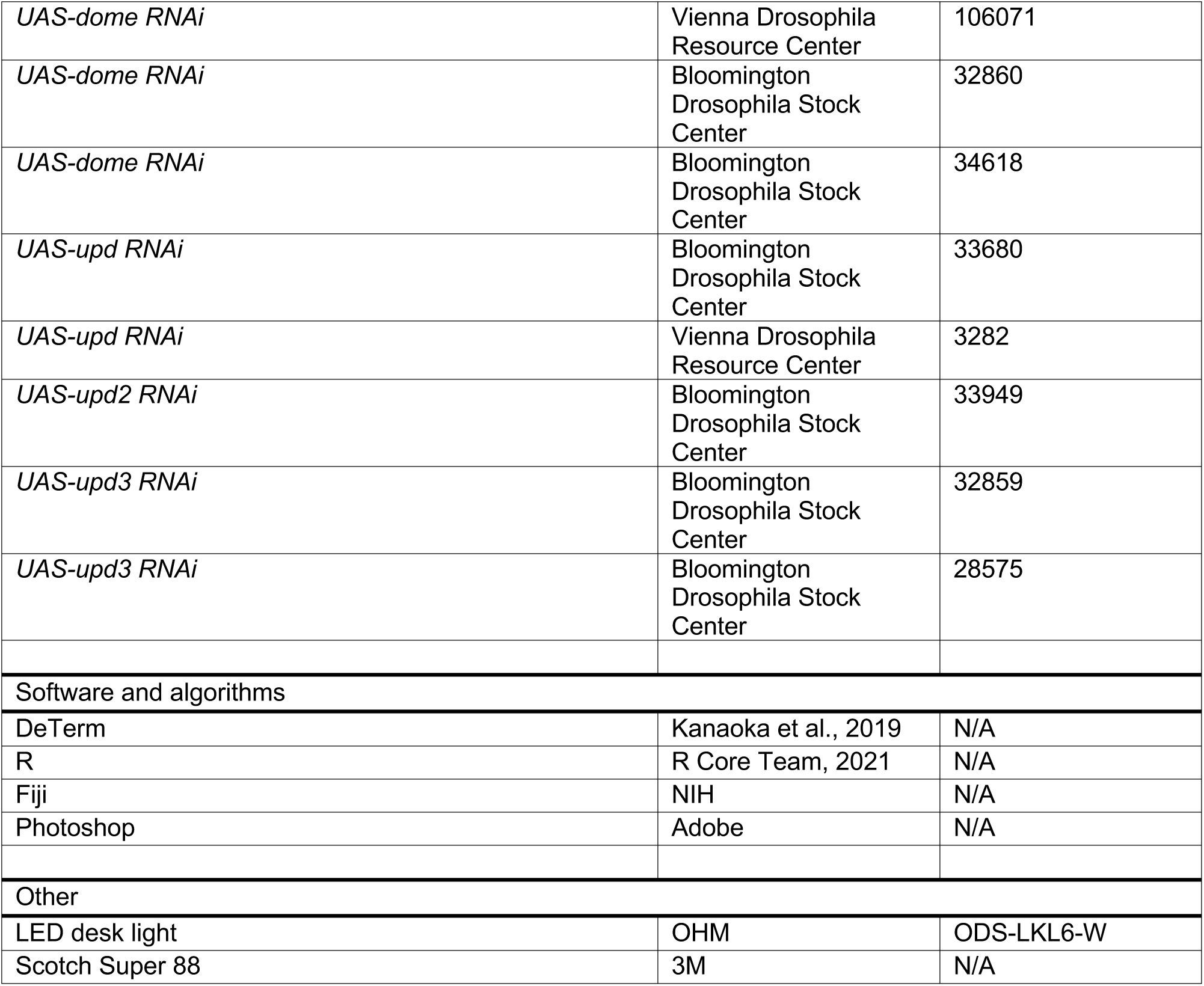
Resource table.

### Experimental diets

We cooked the high yeast diet (HYD) or low yeast diet (LYD) based on semidefined media (SDM) as described previously (Watanabe et al., 2017). The original SDM recipe is described at the Bloomington Drosophila Stock Center <https://bdsc.indiana.edu/information/recipes/germanfood.html>. HYD and LYD were composed of brewer’s yeast (MPBio 2903312), glucose (Wako 049-31165), sucrose (Wako 196-00015), peptone (Fluka 82303), and agar (Matsuki Kanten). The complete compositions of these diets can be found in Supplementary File 1. After the ingredients were mixed, water was added to a final volume of 200 ml, followed by autoclaving. Once the foods had cooled, 1.2 ml propionic acid (Nacalai Tesque, 29018-55) and 2 ml 10% butyl p-hydroxybenzoate (Nacalai Tesque, 06327-02) in 70% ethanol were added. The foods were then dispensed into vials and left overnight before use.

For the supplementation with essential amino acid solution, we used 50x MEM EAA solution (Wako 132-15641). Each fraction of holidic medium (Piper et al., 2014, 2017) other than amino acids (vitamins, cholesterol, metal ion and other ingredients) was added to LYD at 10 times the concentration in holidic medium. Amino acids mixture from holidic medium was added at a 1x or 3x concentration. The complete compositions of diets used in nutrient supplementation experiments can be found in Supplementary File 1.

### Imaging and quantification for assessing dendritic morphology

Images of ddaC, one of the class IV neurons, in A3–A5 segments were acquired in live whole-mount larvae as described (Hattori et al., 2013; Parrish et al., 2009; Matsubara et al., 2011). Protocols for single-cell labeling (MARCM) were as previously described (Shimono et al., 2014). For quantification of the number of dendritic branching terminals, we drew an outline of the dendritic field as a region of interest (ROI) by connecting the outermost dendritic terminals with the Adobe Photoshop path tool. Then, dendritic branching terminals inside the ROI were semi-automatically counted by using DeTerm (Kanaoka et al., 2019). In addition, the area size of the ROI was measured as the dendritic coverage size. Some representative control images and control data are shared by multiple figures. See figure legends.

### Immunostaining

Dissected wandering 3^rd^ instar larvae were fixed in a 1:10 dilution of Formaldehyde Solution (Nacalai Tesque, 16222-65) in PBS plus 0.05 % Triton X-100 for 30 min, then washed 3 times in PBS plus 0.1% Triton X-100 (PBST). After blocking in PBST plus 2% bovine serum albumin for 30 min, primary antibodies listed in Table 1 were added, then incubated overnight at 4°C. After 3 successive washes, secondary antibodies were added, then incubated for 1h at room temperature. Finally, samples were mounted using FluorSave™ Reagent (Calbiochem).

### Quantification of signal intensity

To quantify signal levels in muscle, we made Z-stack images and chose muscle 9, one of the closest muscles to ddaC, for measuring the signal intensity. For quantification of Wg or 10x Stat-GFP signals, we measured the signal intensity inside a 57-pixel or a 40-pixel square ROI, respectively. Three ROIs were drawn for each muscle, and the average value was calculated. For quantification of RedStinger driven by *wg-Gal4*, the signal intensity inside nuclei that were identified by DAPI signals was measured. Then, the values from 2-7 nuclei in each muscle were averaged. For quantification of p-Akt levels in cell bodies of ddaC neurons, we selected the single section containing the strongest signal in ddaC and measured signal intensities inside a 10-pixel square ROI located on the ventral side of ddaC nuclei.

### Electrophysiology

Extracellular single-unit recordings in wandering 3rd instar larvae were performed as previously described (Terada et al., 2016; Onodera et al., 2017). We recorded the activity of v’ada of class IV neurons, which showed hyperarborization on LYD. For blue light irradiation, the 460-495 nm light at 72mW/mm^2^ power was illuminated for five seconds. The light spot was 1.5 mm in diameter. Peristimulus time histograms were calculated at 250-ms bins. The mean spontaneous firing frequencies were quantified in the 20-s window preceding the light stimuli. The mean firing frequencies during the light stimulation were quantified in the 5-s entire window. The firing changes were calculated by subtracting the mean spontaneous firing rate from the light-evoked one (Change amount) or using a ratio of the mean spontaneous firing rate divided by the mean light-evoked one (Change rate).

### Light/dark choice assay

Light/dark choice assays were performed as previously described with modifications (Yamanaka et al., 2013). For the assay of foraging larvae, we used foraging 3^rd^ instar larvae one day before they start wandering. We prepared 2% agar plate with a lid half of which was covered with black tape and 20 larvae were placed along the junction between light and dark sides. After the plates were illuminated for 15 min with white LED light at 700 lux, the number of larvae in both dark and light areas were counted. In some trials, one or two larvae dug into the agar. Such larvae were excluded from calculation of dark preference index. For the assay of wandering larvae, two opposed plastic tubes were joined by transparent scotch tape and one of the vials was covered with black tape. After 16 wandering larvae reared on HYD or LYD were put near the junction of the tubes, they were illuminated by the 700-lux light for 15min, then the number of larvae in both dark and light areas were counted. Dark preference index was calculated as follows:

((Number of larvae in dark) − (Number of larvae in light))⁄(Total number of larvae)

### RNA-seq

Protocols for sample preparation and data analysis of RNA-seq were essentially as described in Watanabe et al. (2019). To prepare each replicate, RNA was extracted from 5 whole bodies of male wandering third-instar larvae. The following procedures are different from Watanabe et al. (2019): 1) the NEBNext Ultra II Directional RNA Library Prep Kit for Illumina (NEB, E7760) was used for library preparation. 2) RNA-sequencing was performed on an Illumina NextSeq 500 system using single end reads. 3) All raw sequencing data were trimmed using TrimGalore (ver. 0.6.0, Cutadapt ver. 1.18; DOI:10.5281/zenodo.5127899, DOI:10.14806/ej.17.1.200) with - clip_R1 13 option. 4) Gene-based read counts were obtained using htseq-count (ver. 0.11.3; Anders et al., 2015) with -s reverse -a 10 options. 5) Differential expression analysis was performed on the count data using a generalized linear model (GLM) in the edgeR Bioconductor package (ver. 3.30.3; McCarthy et al., 2012; Robinson et al., 2010). All the RNA-sequencing data have been deposited and are available in the DDBJ Sequence Read Archive. The accession numbers for the data are DRR311224-DRR311229 (BioProject accession number: PRJDB12048).

### Statistical analysis

R (R Core Team, 2021) was used for stastical analysis. Values of P < 0.05 were considered statistically different. Student’s t-test or the Wilcoxon-Mann-Whitney test was used for two-group comparisons, and Dunnett’s test, Steel test, or Steel-Dwass tests were used for multiple comparisons. We also analyzed interaction effects between genotype and diet using two-way analysis of variance (ANOVA). Statistical tests used, the exact sample size (n), and P values are shown in Supplementary File 3. R was also used to draw 95% confidence ellipses. See also figure legends for details.

## ACKNOWLEDGMENTS

The reagents and genomic datasets were provided by the *Drosophila* Genetic Resource Center at Kyoto Institute of Technology, National Institute of Genetics, the Bloomington Stock Center, Vienna *Drosophila* Resource Center and FlyBase. We thank T. Kondo and Y. Sando for performing RNA-sequencing; J. A. Hejna for polishing the manuscript; T. Kambe, R. Niwa, T. Jovanic, N. Yamanaka, S. Goulas and other members of the Uemura laboratory for discussions and their technical assistance; M. M. Rolls, A. Wodarz, T. Igaki, T. Ito, M. Nakamura, M. Yamazaki and M. Sato for kindly providing reagents; and Y. Xiang for sharing unpublished results. This work was supported by AMED-CREST (JP18gm1110001 to T. Ue), Japan Society for the Promotion of Science (JSPS; 15H02400 to T. Ue., 21H00251 and 21K06186 to Y.H., 20J15084 to Y.K.), JST FOREST Program (JPMJFR2051 to Y.H.), the Naito Foundation (to Y.H.), and the Japan Foundation for Applied Enzymology (to Y.H.).

## AUTHOR CONTRIBUTIONS

Y.H. and T.Ue. conceived and designed the study. Y.K. and Y.H. designed experiments. Y.K. performed most of the experiments. Y.H. analyzed RNA-seq data. K.O. and T.Usui. performed electrophysiological experiments. K.W. performed the RNA-seq experiment. Y.H., Y.K., and T.Ue. wrote and edited the manuscript, with contributions from all authors.

## Figure supplements

**Figure 1—figure supplement 1.**
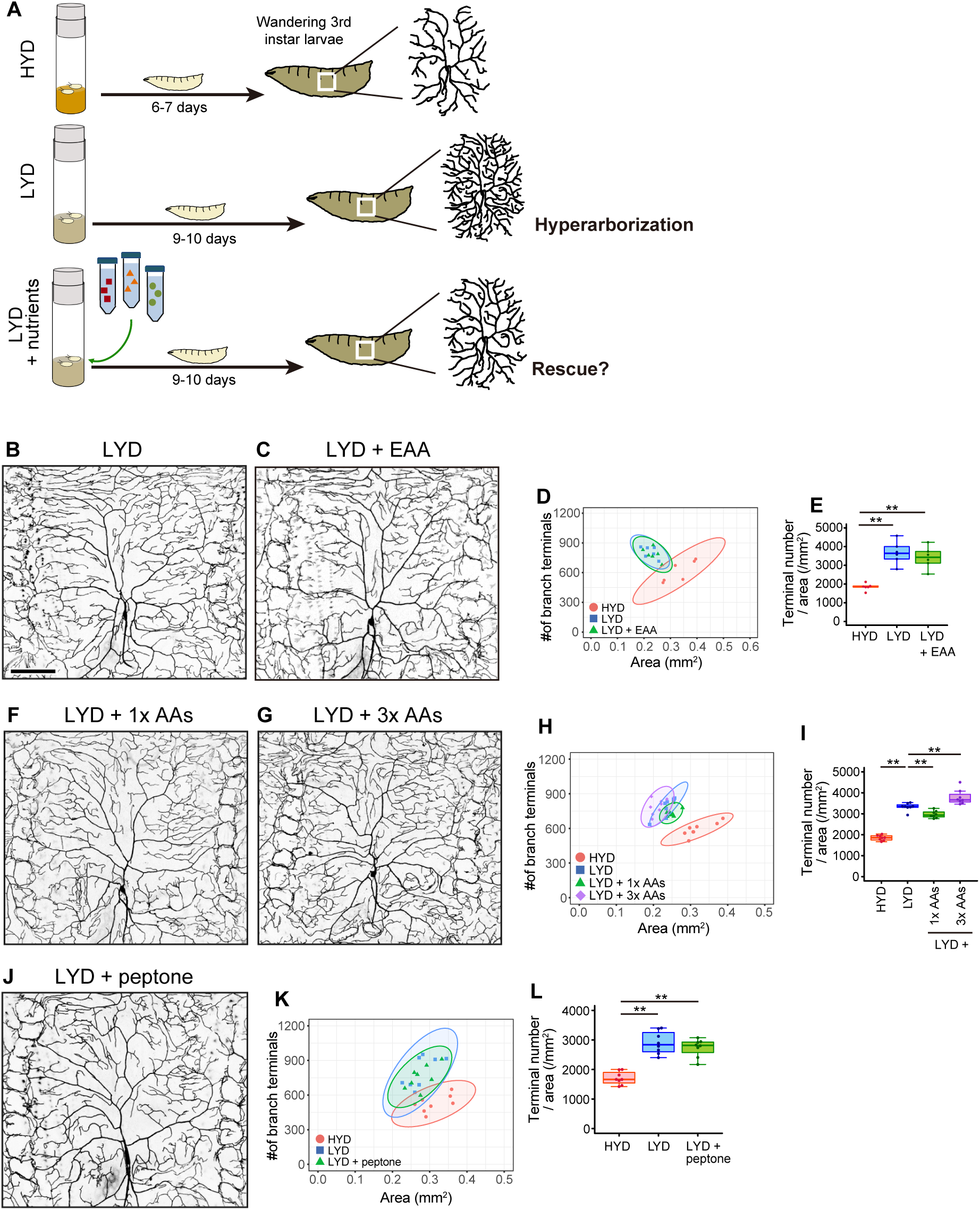
Addition of amino acids does not rescue the hyperarborization. (A) A schematic diagram outlining the observation of class IV neurons in larvae reared on the different diets. We collected wandering 3rd instar larvae and imaged class IV neurons as scheduled throughout our dietary or genetic interventions. This is because the larval development took longer on LYD or LYD + nutrients compared to HYD. (B-E) Images of ddaC neurons in larvae reared on LYD (B) or LYD + essential amino acids (LYD + EAA; C). Two-dimensional plot (D) and densities of branch terminals (E; One-way ANOVA and Tukey’s HSD test, n = 6). The 95% confidence elipse of LYD + EAA almost totally overlaps with that of LYD and remains largely separare from that of HYD (D). (F-I) Images of ddaC neurons in larvae reared on LYD + 1x amino acids, which is the same concentration as in the holidic medium (LYD + 1x AAs; F), or LYD + 3x AAs (G). Two-dimensional plot (H) and densities of branch terminals (I; Steel test, n = 8). Terminal density on LYD + 1x AAs was significantly lower than on LYD, whereas that on LYD + 3x AAs was higher (I). In contrast to these opposoite effects, both the elipse of LYD + 1x AAs and that of LYD + 3x AAs partly overlap with the LYD elipse and both are located apart from that of HYD (H). These effects of supplementing amino acids to LYD make sharp contrast to that of supplementing VMCO or VMC (Figures 1J and 1T). (J-L) Images of ddaC neurons in larvae reared on LYD + peptone (J). Two-dimensional plot (K) and densities of branch terminals (L; Steel test, n = 8). The elipse of LYD + peptone almost totally overlaps with that of LYD (K). Boxplots in (E, I, and L) are depicted as in Figure 1C. **P < 0.01. Scale bar, 100μm.

**Figure 2—figure supplement 1.**
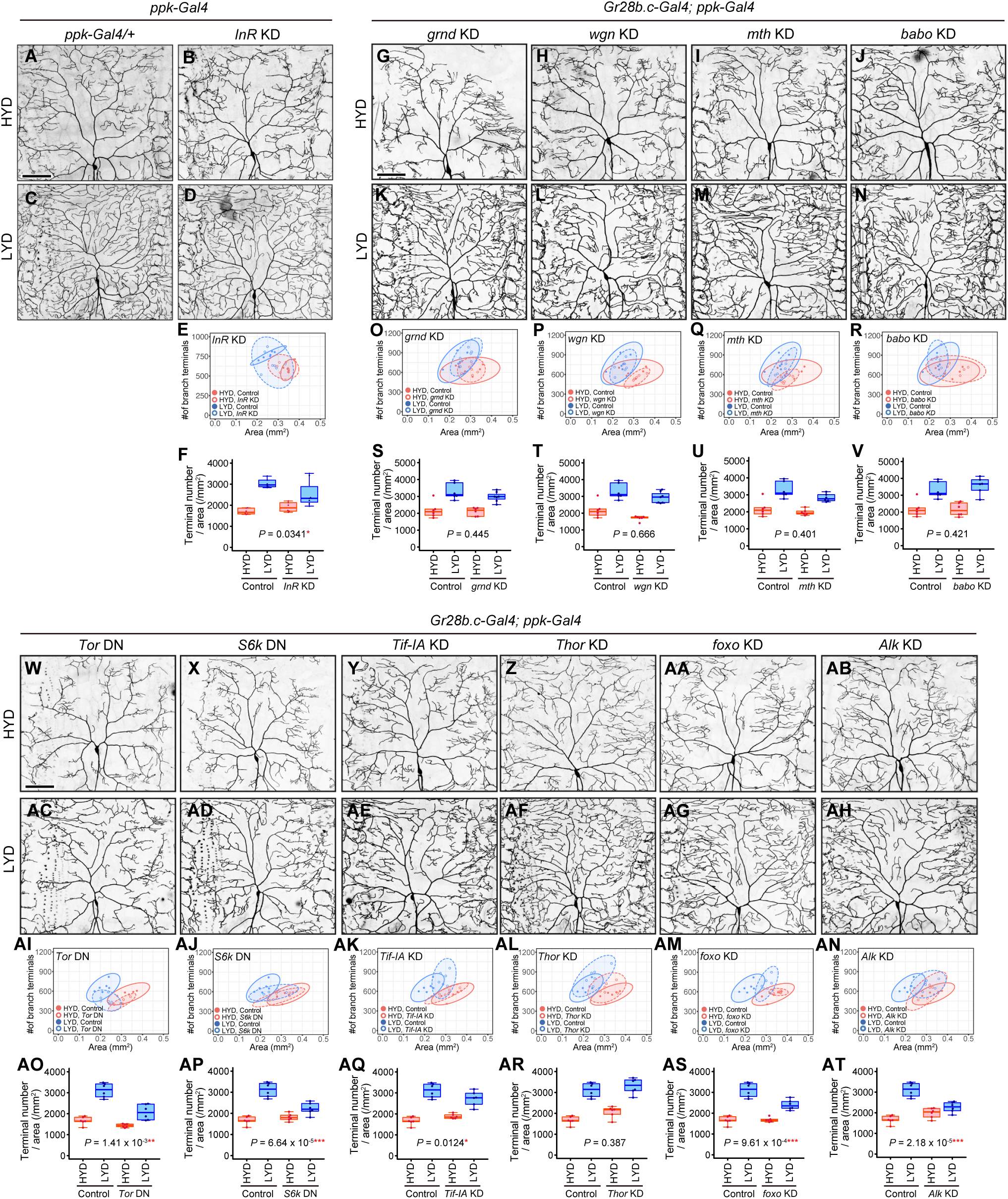
Contributions of intracellular signaling factors or Akt signaling components to the hyperarborization. (A-F) Images of control (A and C) or *InR* knockdown ddaC neurons (B and D), on HYD or LYD. *UAS-InR RNAi ^BL51518^* was used in this experiment. Two-dimensional plot (E) and densities of branch terminals (F). The difference between the diets in *InR* knockdown ddaC neurons is significantly smaller compared to that in control ddaC neurons as indicated by the P-value (Two-way ANOVA, n= 6). We also knocked down *InR* using another *RNAi* line (*UAS-InR RNAi^BL31594^*), but *InR* knockdown ddaC neurons in this line did not significantly ameliorate the hyperarborization (data not shown). (G-V) Images of *gnrd* (G and K), *wgn* (H and L), *mth* (I and M), or *babo* (J and N) knockdown ddaC neurons, on HYD or LYD. Images of control ddaC neurons are shown in Figure 2A and 2D. Two-dimensional plots (O-R) and densities of branch terminals (S-V). The differences between HYD and LYD in control and *gnrd*, *wgn*, *mth*, or *babo* knockdown ddaC neurons are not significantly different as indicated by the respective P-values (Two-way ANOVA, n= 6). (W-AT) Images of *Tor* (W and AC), *S6k* (X and AD), *Tif-IA* (Y and AE), *Thor* (Z and AF), *foxo* (AA and AG), or *Alk* (AB and AH) knockdown ddaC neurons, on HYD or LYD. Two-dimensional plots (AI-AN) and densities of branch terminals (AO-AT). The differences between HYD and LYD in control and *Tor*, *S6k*, *Tif-IA*, *foxo*, or *Alk* knockdown ddaC neurons are significantly different as indicated by the respective P-values (Two-way ANOVA, n= 6). We also knocked down *Alk* using *ppk-GAL4*, but *Alk* knockdown ddaC neurons did not show significantly ameliorated hyperarborization (data not shown). Control data in (O-R) and (S-V) are shared with Figures 2G and 2H and those in (AI-AN) and (AO-AT) are shared with Figures 2I and 2J. Boxplots in (F, S-V and AO-AT) are depicted as in Figure 1C. *P < 0.05, **P < 0.01, and ***P < 0.001. Scale bars, 100μm.

**Figure 3—figure supplement 1.**
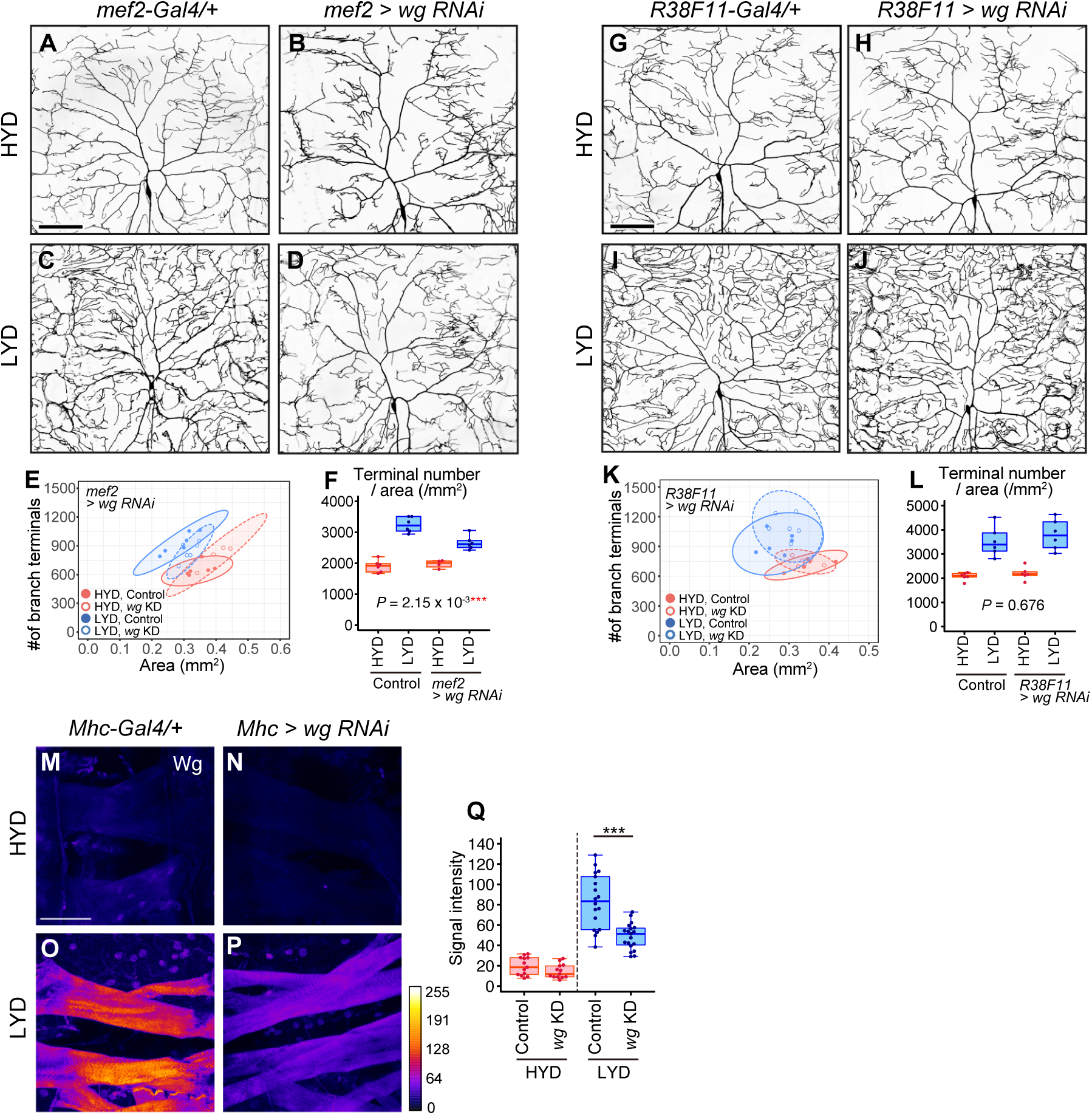
Wg from muscles, but not from epidermal cells, contributes to the hyperarboriztion phenotype. (A-F) Images of ddaC neurons in control larvae (A and C) or larvae with *wg* knocked down in muscles using *mef2-Gal4* (B and D), on HYD or LYD. Two-dimensional plot (E) and densities of branch terminals (F). The differences between HYD and LYD in control larvae and larvae with *wg* knocked down in muscles are significantly different, as indicated by the P-value (Two-way ANOVA, n= 4-6). (G-L) Images of ddaC neurons in control larvae (G and I) or larvae with *wg* knocked down in epidermal cells (H and J), on HYD or LYD. Two-dimensional plot (K) and densities of branch terminals (L). The differences between HYD and LYD in control larvae and larvae with *wg* knocked down in epidermal cells are not significantly different as indicated by the P-value (Two-way ANOVA, n= 6). (M-Q) Muscles in control larvae (M and O) or larvae with *wg* knocked down in muscles using *Mhc-gal4* (N and P), on HYD or LYD, were stained for Wg. The signal intensities correspond to the indicated color code. (Q) Quantification of the mean Wg immunofluorescence in muscle 9 (Wilcoxon-Mann-Whitney test, n = 13-19). Knocking down *wg* decreased the signal intensity in the muscle on LYD. Boxplots in (F, L and Q) are depicted as in Figure 1C. ***P < 0.001. Scale bars, 100μm.

**Figure 4—figure supplement 1.**
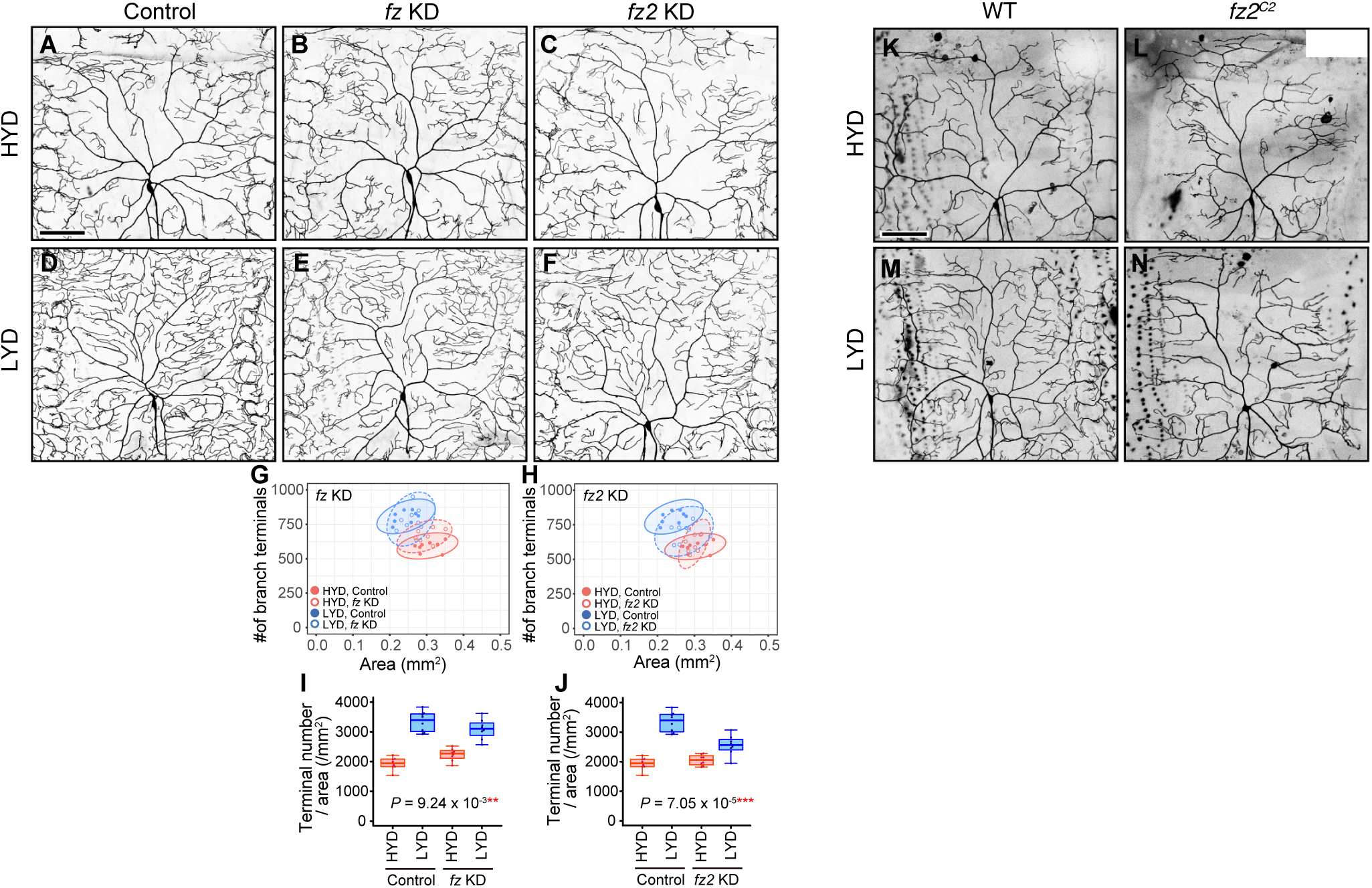
Fz2, a receptor for Wnt proteins, is required in class IV neurons to hyperarborize their dendrites. (A-J) Images of control (A and D), *fz* knockdown (B and E), or *fz2* knockdown (C and F) ddaC neurons, on HYD or LYD. Two-dimensional plots (G and H) and densities of branch terminals (I and J). The differences between HYD and LYD in control and *fz* or *fz2* knockdown ddaC neurons are significantly different, as indicated by the respective P-values (Two-way ANOVA, n= 8). (K-N) Images of ddaC neurons from a wild-type clone (WT; K and M) or a *fz2^C2^* mutant clone (L and N), on HYD or LYD. Control data in (G) and (I) are shared with (H) and (J). Boxplots in (I) and (J) are depicted as in Figure 1C. **P < 0.01, and ***P < 0.001. Scale bars, 100μm.

**Figure 4—figure supplement 2.**
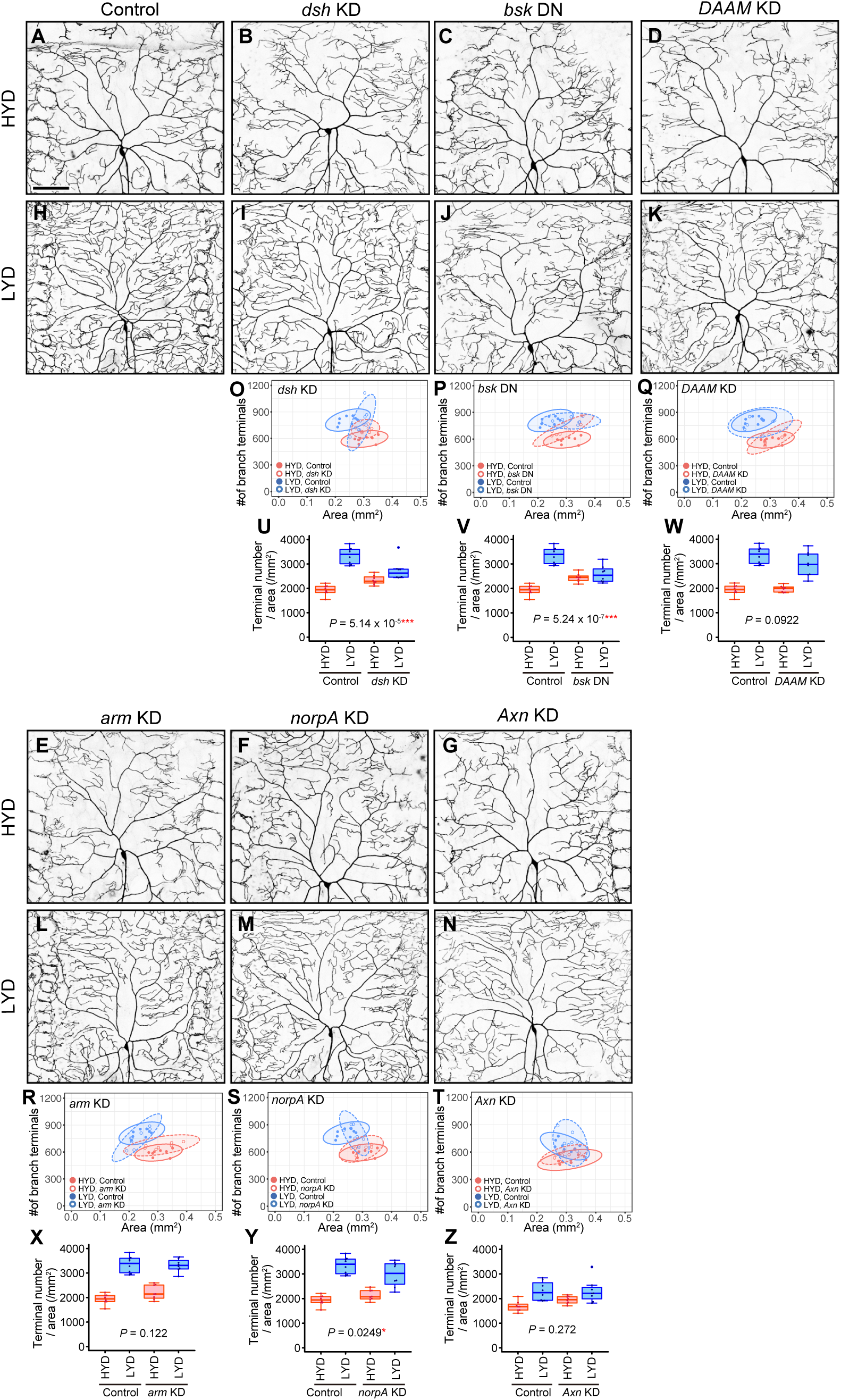
Effects of inhibiting intracellular Wnt signaling components on hyperarborization. (A-N) Images of control ddaC neurons (A and H), *dsh* knockdown ddaC neurons (B and I), ddaC neurons expressing a dominant-negative form of *basket* (*bsk DN*) (C and J), *DAAM* knockdown ddaC neurons (D and K), *arm* knockdown ddaC neurons (E and L), *norpA* knockdown ddaC neurons (F and M), or *Axin* knockdown ddaC neurons (G and N), on HYD or LYD. (O-Z) Quantitative analysis. (O-T) Two-dimensional plots. (U-Z) Densities of branch terminals. The difference between HYD and LYD in *dsh* knockdown (U), *bsk DN* expressing (V), or *norpA* knockdown (Y) ddaC neurons is significantly different from that in control ddaC neurons, as indicated by the respective P-values (Two-way ANOVA, n= 8). However, knocking down *dsh* or blocking JNK signaling tended to increase the branch density over the control genotype on HYD, which may consequently reduce the differences in densities of dendritic terminals between the diets. The effect of *norpA* knockdown was marginal and the distribution of the terminal densities was largely unaffected. Images of control neuron (A and H) are shared with Figure 4—figure supplement 1A and B. Control data in (O-S) and (U-Y) are shared with Figure 4—figure supplement 1G and 1I. Boxplots in (U-Z) are depicted as in Figure 1C. *P < 0.05, and ***P < 0.001. Scale bars, 100μm.

**Figure 5—figure supplement 1.**
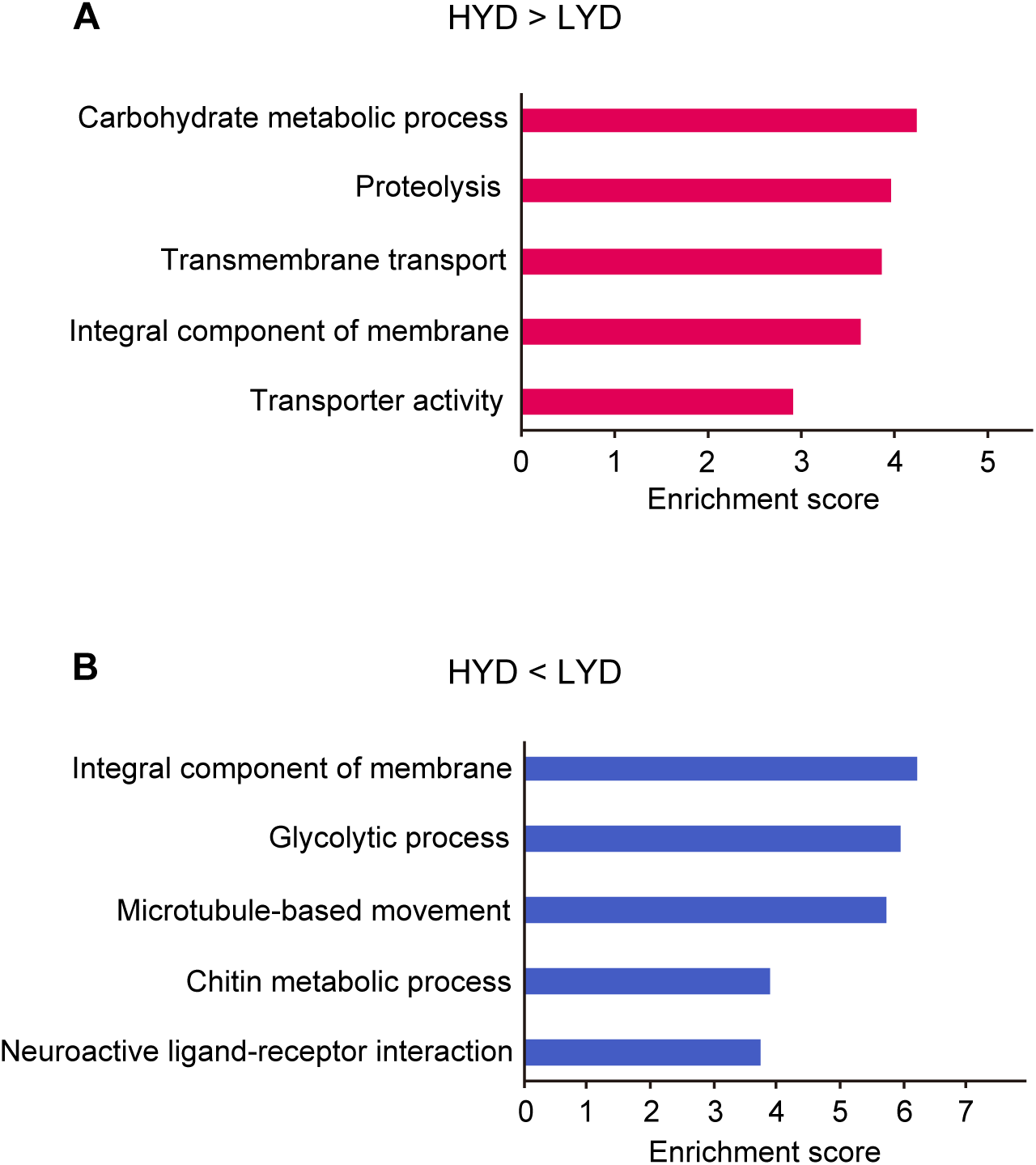
Enriched terms in functional annotation clustering of differentially expressed genes depending on diets in whole body RNA-seq data. Enrichment score of the top five functional annotation clusters enriched in genes highly expressed on HYD rather than on LYD (A) or genes highly expressed on LYD rather than on HYD (B) in whole body RNA-seq data.

**Figure 5—figure supplement 2.**
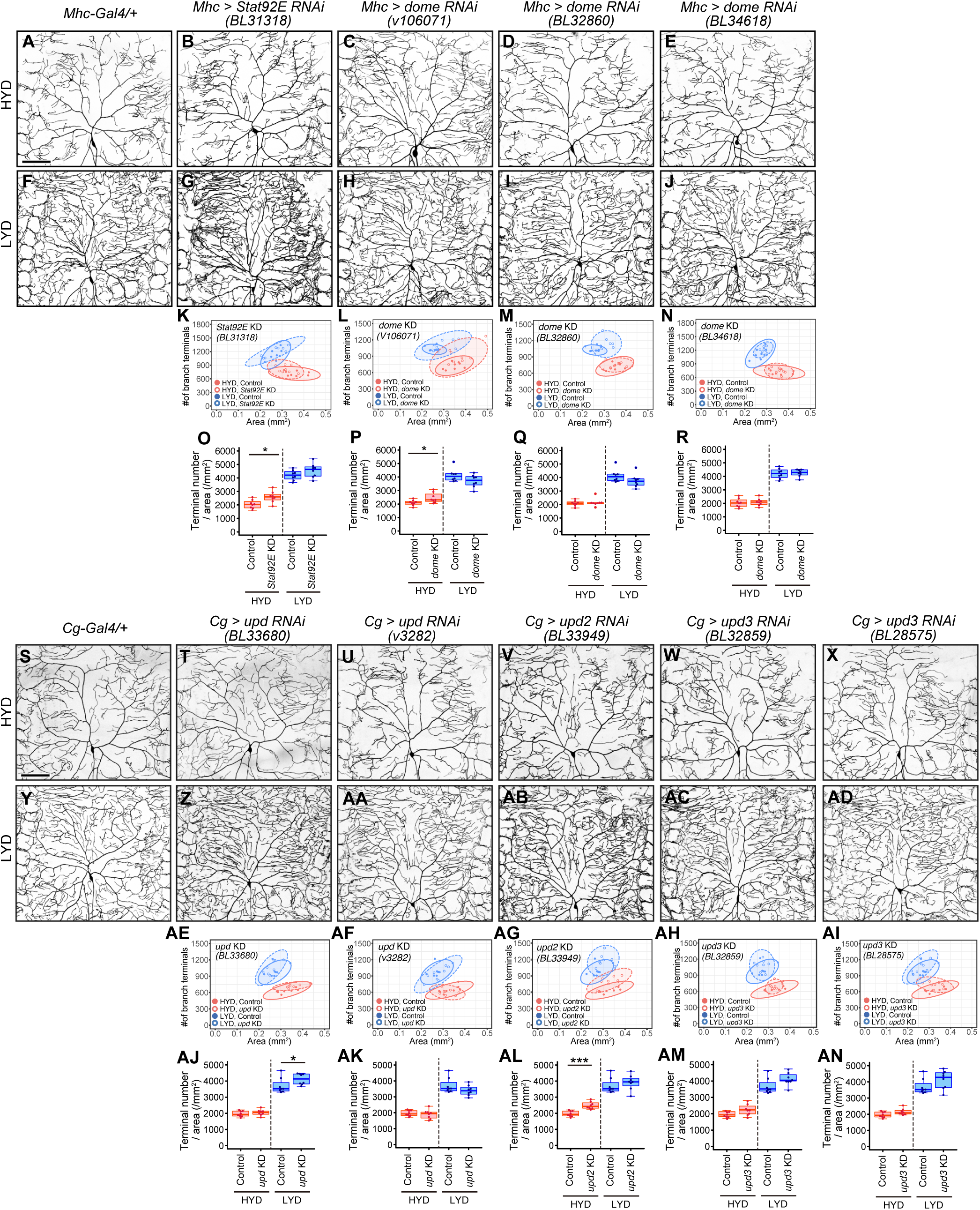
Effects of inhibiting components of JAK/STAT pathway on hyperarborization. (A-R) Images of ddaC neurons in control larvae (A and F) and in larvae with *Stat92E* (B and G) or *dome* (C-E and H-J) knocked down in muscles, on HYD or LYD. Two-dimensional plots (K-N) and densities of branch terminals (O-R). (S-AN) Images of ddaC neurons in control larvae (S and Y) and in larvae with *upd* (T, U, Z and AA), *upd2* (V and AB), or *upd3* (W, X, AC and AD) knocked down in the fat body and hemocytes, on HYD or LYD. Two-dimensional plots (AE-AI) and densities of branch terminals (AJ-AN). Images of control neuron (S and Y) are shared with Figure 5AA and 5AC. Control data in (K) and (O) are shared with (N) and (R), those in (L and M) and (P and Q) are shared with Figures 5Q and 5S, and those in (AE-AI) and (AJ-AN) are shared with Figures 5AE and 5AF. Boxplots in (O-R and AJ-AN) are depicted as in Figure 1C. *P < 0.05, and ***P < 0.001 (Wilcoxon-Mann-Whitney test, n =8). Scale bars, 100μm.

## Supplementary Files

**Supplementary File 1:**

**Compositions of the experimental diets.**

**Supplementary File 2:**

**RNA-seq data of larval whole bodies at the wandering third-instar stage on HYD or LYD.**

(A) List of Differentially expressed genes between HYD and LYD in whole larval bodies at the wandering third-instar stage (adjusted P value < 0.05).

(B and C) List of functional annotation clusters that significantly enriched (enrichment score ≥ 1.3) in genes highly expressed on HYD rather than on LYD (B) or genes highly expressed on LYD rather than on HYD (C).

**Supplementary File 3:**

**Statistical details of experiments and a list of genotypes.**

